# Spatial analysis of NOS2 and COX2 interaction with T-effector cells reveals immunosuppressive landscapes associated with poor outcome in ER- breast cancer patients

**DOI:** 10.1101/2023.12.21.572867

**Authors:** Lisa A. Ridnour, Robert Y.S. Cheng, William F. Heinz, Milind Pore, Ana L. Gonzalez, Elise L. Femino, Rebecca Moffat, Adelaide L. Wink, Fatima Imtiaz, Leandro Coutinho, Donna Butcher, Elijah F. Edmondson, M. Cristina Rangel, Stephen T.C. Wong, Stanley Lipkowitz, Sharon Glynn, Michael P. Vitek, Daniel W. McVicar, Xiaoxian Li, Stephen K. Anderson, Nazareno Paolocci, Stephen M. Hewitt, Stefan Ambs, Timothy R. Billiar, Jenny C. Chang, Stephen J. Lockett, David A. Wink

**Author notes:** Primary Author(s) (Lisa A. Ridnour, Robert Y.S. Cheng, William F. Heinz, Milind Pore).

## Abstract

Multiple immunosuppressive mechanisms exist in the tumor microenvironment that drive poor outcomes and decrease treatment efficacy. The co-expression of NOS2 and COX2 is a strong predictor of poor prognosis in ER- breast cancer and other malignancies. Together, they generate pro-oncogenic signals that drive metastasis, drug resistance, cancer stemness, and immune suppression. Using an ER- breast cancer patient cohort, we found that the spatial expression patterns of NOS2 and COX2 with CD3+CD8+PD1- T effector (Teff) cells formed a tumor immune landscape that correlated with poor outcome. NOS2 was primarily associated with the tumor-immune interface, whereas COX2 was associated with immune desert regions of the tumor lacking Teff cells. A higher ratio of NOS2 or COX2 to Teff was highly correlated with poor outcomes. Spatial analysis revealed that regional clustering of NOS2 and COX2 was associated with stromal-restricted Teff, while only COX2 was predominant in immune deserts. Examination of other immunosuppressive elements, such as PDL1/PD1, Treg, B7H4, and IDO1, revealed that PDL1/PD1, Treg, and IDO1 were primarily associated with restricted Teff, whereas B7H4 and COX2 were found in tumor immune deserts. Regardless of the survival outcome, other leukocytes, such as CD4 T cells and macrophages, were primarily in stromal lymphoid aggregates. Finally, in a 4T1 model, COX2 inhibition led to a massive cell infiltration, thus validating the hypothesis that COX2 is an essential component of the Teff exclusion process and, thus, tumor evasion. Our study indicates that NOS2/COX2 expression plays a central role in tumor immunosuppression. Our findings indicate that new strategies combining clinically available NOS2/COX2 inhibitors with various forms of immune therapy may open a new avenue for the treatment of aggressive ER- breast cancers.

## Introduction

Immune therapies, such as checkpoint inhibitors, cancer vaccines, and T cell adoptive therapies, have created novel treatment options for advanced cancers (1–6). Some success in combining immune with conventional therapy has shown immune system polarization as a significant factor in overcoming clinically challenging tumors. Various immunosuppressive mechanisms within the tumor microenvironment (TME) are among the most significant stumbling blocks for successfully treating challenging cancers (7). Given the importance of immune evasion for tumor survival, the spatial characterization of immune cells and the distribution of potential immunosuppressive agents within the TME must be evaluated.

The numerous molecular effectors inhibiting the immune response within the TME act on the immune, stromal, and tumor cells compartments. Immunosuppressive mechanisms include T cell-centered mechanisms, such as PD1/PDL1 and Treg; cellular factors, cytokines, growth factors, and small molecules derived from metabolites, such as kynurenines and polyamines (8–11). The co-existence of these different immunosuppressive mechanisms in the TME provides multiple layers of immune suppression.

Nitric oxide synthase 2 (inducible NOS; NOS2) and cyclooxygenase-2 (COX2) have been suggested as drivers of a poor prognosis (12–17), and play a significant role in the immunosuppressive properties of several cancers. In many solid tumors, NOS2 and COX2 are elevated (18–23). In estrogen receptor negative (ER-) breast cancer, in particular, elevated NOS2/COX2 levels are one of the strongest predictors of poor prognosis (16). Both NOS2 and COX2 form a feedforward loop fueling the propagation of different pro-tumorigenic mechanisms. They, in turn, activate primary oncogenic pathways, ultimately leading to metastasis, chemoresistance, and cancer stem cell characteristics (16). According to recent research (12, 24), NOS2 and COX2 are associated with the restriction of CD8 cells in human malignancies. Moreover, in animal models, inhibiting NOS2 and COX2 produces profound changes in the immune system, countering the metastatic process and tumor growth, resulting in cures and resistance to tumor rechallenge (24). Accordingly, NOS and COX inhibitors dramatically increase survival in chemo-resistant triple-negative breast cancer (TNBC) (24–26).

Mechanistically, NOS2 and COX2 induce the cytokines IL10, TGFβ, and IL6, which suppress the immune system (24, 27), while modifying the metabolic programming of particular cells (16, 28–30). NOS2 and COX2 are heme proteins that can swiftly respond to environmental changes, such as hypoxia and inflammation, making them crucial components linking environmental sensing and immune status (31). CD8 T cells and IFNγ are essential for NOS2 and COX2 expression in the tumor cells *in vivo* (12). This requirement of antitumor factors to induce NOS2 and COX2 is strongly associated with poor outcomes and presents a conundrum. Indeed, regions of elevated NOS2 and COX2 are close to restricted CD8 cells and lymphoid aggregates, whereas, in cold regions (i.e., low CD8 counts) and immune deserts, COX2 is present (24). However, when CD8 cells penetrate the tumor nest, NOS2 and COX2 levels in the entire tumor are low, suggesting their significant role in CD8 restriction and exclusion. This progression from an inflammatory region to an immune desert suggests that, in addition to NOS2 and COX2, other immune-suppressive factors account for the transition from inflammation to an immune desert status.

IFNγ and cytokines are essential for tumor eradication as well as the induction of NOS2 and COX2, that is linked to poor prognoses (12, 25). This apparent dichotomy of IFNγ activity between tumor eradication (via facilitated positive immune responses in the host) and NOS2/COX2- associated tumor evasion (via PDL1, T_reg_, B7H4, and IDO1), implicates it as a mediator in a delicate balance (16, 32, 33). Similar dichotomies exist for the immunosuppressive agents IDO1 and PDL1 that are stimulated by IFNγ (34, 35). This negative feedback response to IFNγ results in suppression of the cytolytic activity of T cells. The spatial evaluation of immunosuppressive factors and their relationships can be exploited for more effective treatments against difficult-to-treat forms of cancer.

In the current study, we performed immunohistochemical analyses on ER- breast cancer biopsies to determine whether the spatial distribution of NOS2 and COX2 contributes to other immune suppressive mechanisms involving PD1/PDL1, T_reg_, IDO1, and B7H4 to suppress T_eff_ (CD3CD8PD1^-^) cell function. Our collected evidence demonstrates that NOS2 and COX2 play a crucial and complementary role in the tumor’s immune suppression mechanisms, and that the spatial orientation of immune and tumor cells creates a unique tumor landscape that drives the formation of immune deserts and immune cell restriction, fueling immune suppression, thus tumor progression.

## Results

### Comparison of immunosuppressive markers with patient survival

The composition of immunosuppressive factors in the TME influences the immune profile determining therapeutic response (36–39). There are four distinct methods of immunosuppression: 1) cell-to-cell contact with specific receptors such as PDL1/PD1 and B7H4; 2) generation of diffusible proteins like cytokines and growth factors, such as IL10 and TGFβ; 3) production of small diffusible molecules like metabolic substrates, such as kynurenines, polyamine; 4) small molecule regulators, such as NO and PGE2 (40–48). Thus, each type has a specific spatial distribution to mediate immunosuppression that ranges from direct cell contact agents to those that diffuse throughout the tumor, and even systemically, impacting different tissues throughout the body. To define the immunosuppression mechanisms of NOS2 and COX2 within the tumor, multiplex immunofluorescence was used to define their location. Tumor samples from 21 ER- breast cancer patients (ER-), which included deceased (n=11) and alive (n=10), were imaged (see Materials and Methods). Multiple immune cell markers were evaluated, such as: CD68 for macrophage; T cell markers, CD3, CD4, CD8; tumor markers CK-SOX10; immune modulatory factors, PD1, PDL1, FOXP3, IFNγ, NOS2, and COX2. The percentage of cells containing these markers were then compared with survival outcome (Fig.1A). Only NOS2 and COX2 were statistically associated with a negative outcome as a univariate (Fig.1A). NOS2 was evaluated at 3 levels determined by average cell intensity, to be weak (NOS2w), medium (NOS2m), or strong (NOS2s), showing that increased NOS2 expression correlated with poor outcome (Suppl. Fig. 1A). In contrast, IFNγ demonstrated slightly elevated levels that were marginally associated with a positive outcome. In contrast, the other markers lacked significance (Fig.1A). These results indicate that NOS2/COX2 were the only factors associated with poor prognosis at the single cell level, corroborating previous IHC-based studies (49, 50).

**Figure 1.**
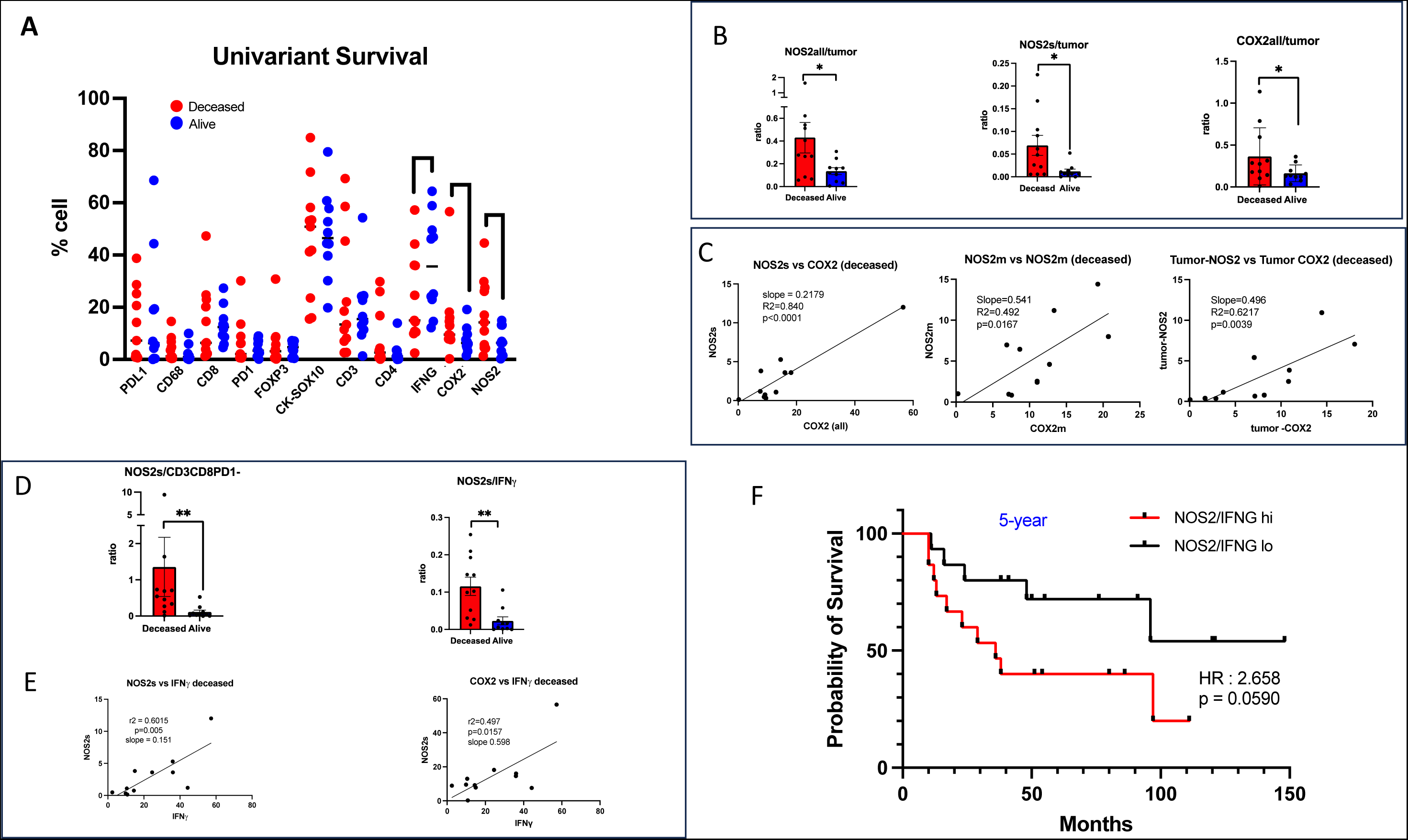
Survival Analysis of Univariant and NOS2 and COX2 with different factors. Comparison of % cell with NOS2 and COX2 with immune markers with respect to survival in ER- breast cancer with deceased n=11 and alive n=10. A) Univariant analysis of the NOS2 and COX2 with PDL1, CD68, CD8, PD1, FOXP3, CK-SOX10, CD3, CD4, IFNγ, COX2, and NOS2 all. B) Ratio of NOS2all (above lowest threshold) (medium deceased = 0.27 : alive = 0.16), NOS2s (medium deceased = 0.048 : alive = 0.0067), and COX2all with tumor (CK-SOX10) (medium deceased = 0.365 : alive = 0.167). C) Linear regression analysis of NOS2 and COX2. D) Ratio of NOS2s with CD3CD8PD1- (T_eff_) (medium deceased=0.54: alive=0.053) and IFNγ (medium deceased= 0.102 : alive=0.0109). E) Linear regression analysis of NOS2s or COX2 with IFNγ. F) Kaplan-Meir plot comparing the ratios of NOS2/CD8A-high and NOS2/CD8A-low; cohort (TCGA-BRCA) divided at median of gene expression ratio (*p <0.5, **p <0.01, *** p<0.001, Mann Whitney test, one-tail).

In the tumor epithelium of ER-breast cancer, there is an inverse relationship between NOS2/COX2 levels and T_eff_ numbers that is associated with poor outcomes. To define the survival relationship between NOS2 and COX2 with other immune and tumor factors, the ratio of the percentage of cells expressing a given factor to NOS2s and COX2 expression levels was evaluated. Prior studies showed that NOS2 is predominantly associated with the tumor epithelium (49). When the ratio of NOS2/COX2 to the tumor marker CK-SOX10 was analyzed (**Fig. 1B**), there was a strong and statistically significant difference in survival, with a significant correlation between higher NOS2/COX2 per tumor cell and decreased survival (**Fig.1B, Suppl. Fig. 1B**). This evidence indicates that higher proportion of NOS2 or COX2 expressing tumor cells is associated with a poor prognosis. Furthermore, a linear analysis showed that NOS2s and COX2 had a strong correlation in deceased but not in alive patients (**Fig. 1C, Suppl.** **Fig.1D**). The analysis of the co-expression of NOS2m and COX2 with the tumor marker CK-SOX10 revealed that tumor-NOS2m and tumor- COX2 was also linear in deceased but not alive patients (**Fig.1C**). Thus, this set of data indicates that tumor-NOS2 and -COX2 correlate strongly with the outcome, confirming our previous finding by immunohistochemistry DAB staining that NOS2 and COX2 in the tumor epithelium is associated with poor outcome (49).

Next, the ratio of NOS2s and COX2 was compared with different immune factors. The results show that the immune markers CD8 and CD3 correlated with survival, while PDL1, PD1, FOXP3, and CD68 did not (**Sup.** Fig.1E**-H**). Further classification of the CD3/CD8 phenotype showed that the ratio of NOS2s and COX2 to CD3+CD8+ (CD3CD8) numbers was significant (**Suppl.** **Fig.1K**). Further subtyping the CD3CD8 phenotypes show that the NOS2s ratio with T_eff_ (CD3CD8PD1^-^) were strongly significant but not with T_ex_ (CD3CD8PD1^+^) (**Fig.1D**, **Suppl. Fig,1K**). Conversely, CD4/CD3 phenotypes such as T_reg_ did not correlate with survival (**Suppl. Fig 1F**). These findings demonstrate that T_eff_ (CD3CD8PD1^-^) has an inverse relationship with NOS2 and COX2. CD8 T_eff_ is a significant source of IFNγ, which can lead to induction of NOS2 and COX2 (12) (Graphical Abstract). Comparing the ratio between NOS2 or COX2 with IFNγ, a strikingly similar significant trend to CD8 T_eff_ was observed (Fig. 1D). Furthermore, the ratios between NOS2, COX2, and CD8 was analyzed using more extensive databases such as GEO and TCGA (**Fig.1F**). Comparing the 75th percentile of high NOS2 to the remaining samples reveals a significant correlation with a negative outcome, confirming the relationship between NOS2 and CD8 (Fig.1F). Therefore, these single-cell fluorescence analyses were confirmed by the larger genomic databases corroborating the existence of a crucial antagonistic relationship between NOS2 and COX2 and T-effector/ IFNγ presence that is associated with survival.

### Immune suppression of Teff cells through negative feedback by NOS2 and COX2

One of the contradictions in this relationship between T-effector/IFNγ and NOS2/COX2 is that IFNγ is required to induce NOS2 and the Th1 cytokines TNF and IL1 in human malignancies (12, 51–53). This evidence suggests that, although NOS2/COX2 inhibit T_eff_/ IFNγ, this cytokine must be present in the tumor. Using a linear analysis, NOS2 and COX2 were plotted against IFNγ in all the samples, showing no significant correlations (**Suppl. Fig. 1I and J**). However, when analyzed comparing deceased and alive patients, the deceased group showed a strong linear relationship between NOS2s and IFNγ, which was absent in the surviving patients . Linear analysis of NOS2 with T_eff_ revealed similar results (**Fig. 1, Suppl. Fig. 1K**). These linear correlations indicate that the NOS2s phenotype requires IFNγ from T_eff_, which leads to increased NOS2/COX2 associated with poor outcomes (paper 4 submitted). The contradicting results between beneficial T_eff_/IFNγ and harmful NOS/COX2 effects can be explained by the ability of PGE2 and NO to decrease Th1 responses via multiple mechanisms, including inducing IL10 and TGFβ (54, 55), in a negative response to IFNγ and TNF/IL1. These survival-based correlations demonstrate that NOS2 and COX2 play a leading and conspicuous role in the immune suppression of Teff cells in ER- breast cancer. Nonetheless, the dichotomy poses the question of how this distribution and configuration may spatially and temporally influence immunosuppression in ER- breast cancer and the relationship between other immune suppressive mechanisms.

### Immune suppressive phenotypes distribution at the whole tumor level

In deceased patients, T_eff_ (CD3CD8PD1^-^) are excluded from tumor nests, while IFNγ is more evenly distributed throughout the tumor (12, 56). To define the spatial localization of T_eff_ and IFNγ relative to NOS2/COX2, density heat maps were utilized to evaluate the juxtaposition between these factors within the TME. Examining the spatial relationship between NOS2 and COX2 with T_eff_ (CD3CD8PD1^-^) and IFNγ reveals their spatially unique orientation (Fig. 2A). In contrast to widely distributed IFNγ throughout the tumor, dense regions of T_eff_ (CD3CD8PD1^-^) clusters form foci spatially distinct from high density NOS2 and COX2 areas (12). While orthogonal, the T_eff_ cells were proximal to NOS2+ regions (Fig. 2A). These findings suggest that in deceased patients, T_eff_ (CD3CD8PD1^-^) are excluded from tumor nests while IFNγ is more evenly distributed throughout the tumor. This is consistent with IFNγ binding to other cells in regions of the lipid rafts (57). From this analysis, there are clear distinct regions of NOS2, COX2, and T_eff_ in the deceased patients.

**Figure 2.**
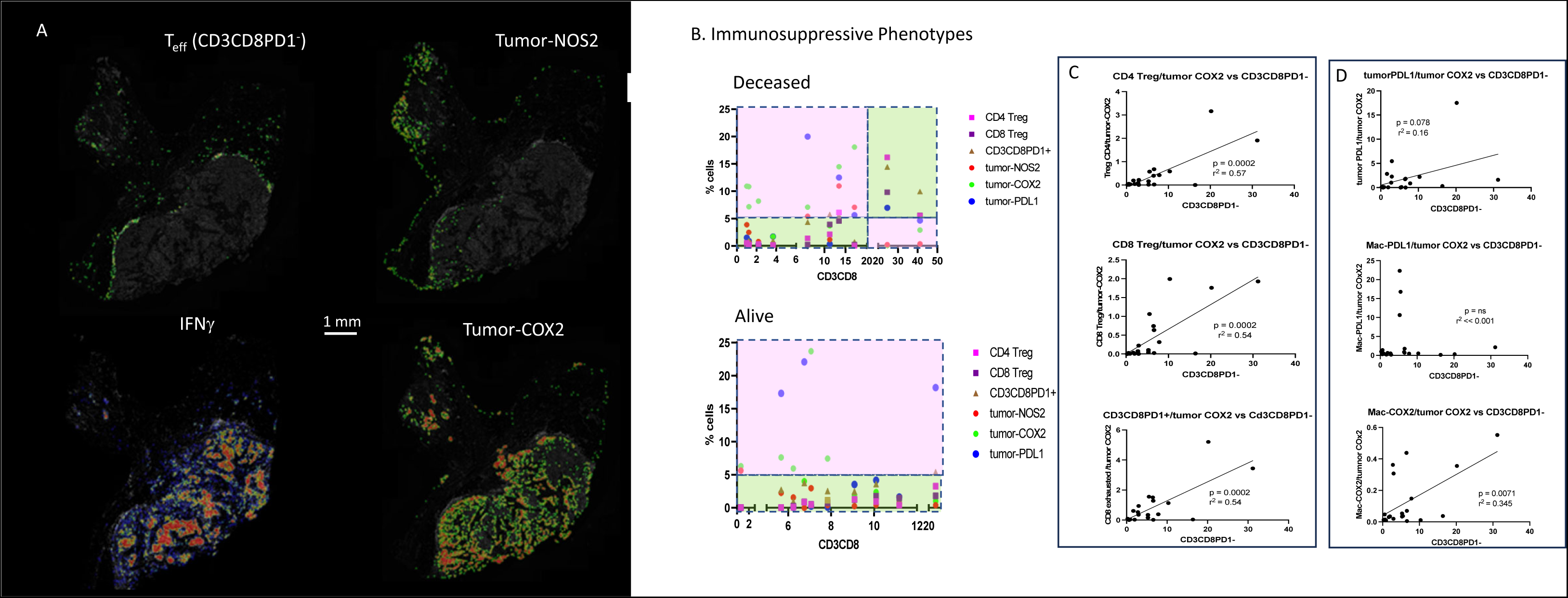
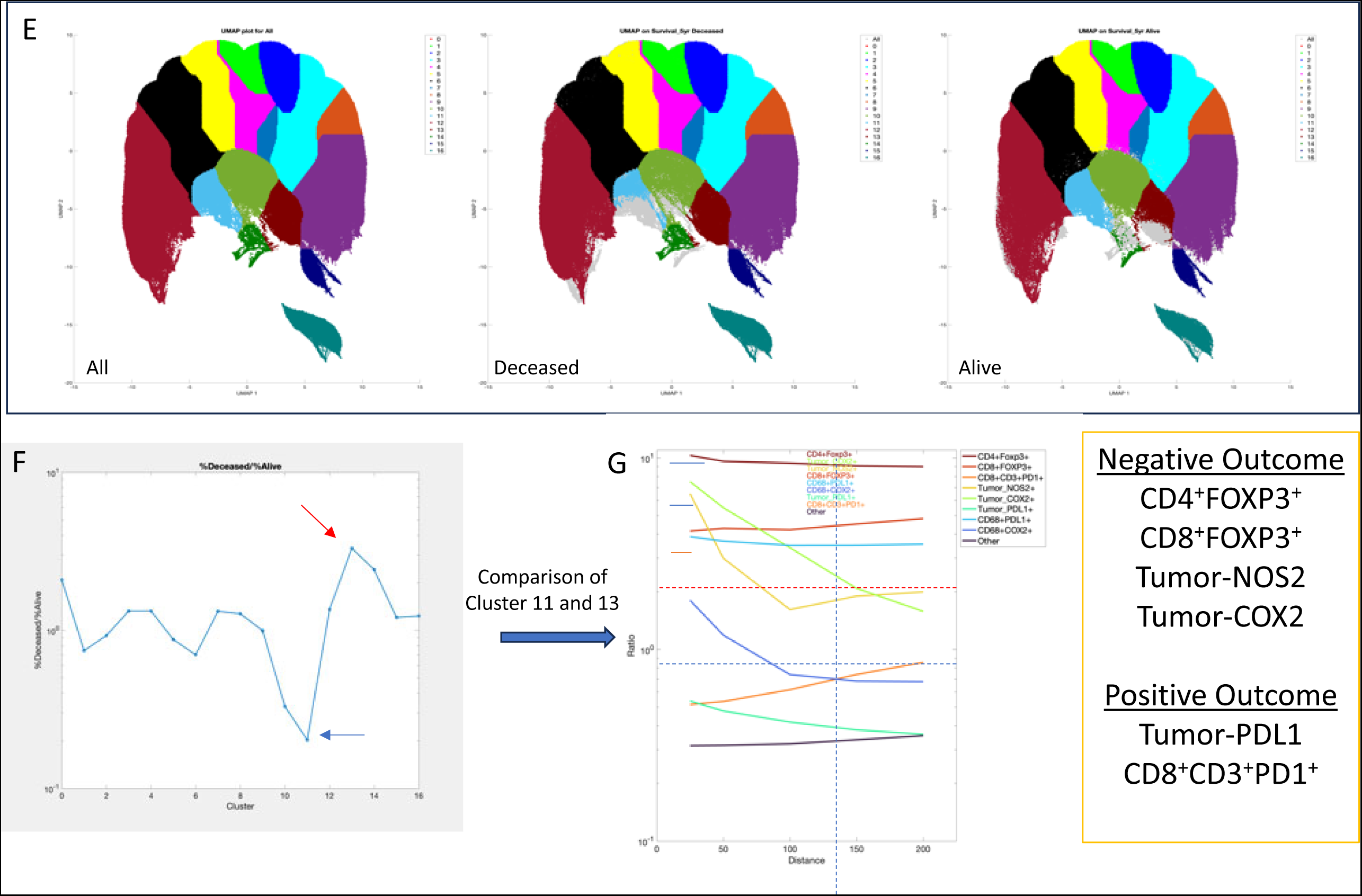
Spatial distribution analysis of NOS2 COX2 with immune cellular phenotypes. A) Density Heat Maps comparing spatial orientation of T_eff_, IFNγ, tumor-NOS2 and tumor-COX2 in representative ER- breast cancer deceased patient. B) Distribution analysis of % cells compared to the CD3CD8 with different tumor and immune cellular phenotypes in Deceased and Alive. C) Linear regression analysis of T cell phenotypes with tumor-COX2 D) linear regression of tumor and macrophage phenotypes with CD3CD8PD1. E) Spatial S-UMAP of the clustering cellular phenotypes. F) cluster analysis comparing survival G) nearest neighborhoods analysis to identify cellular phenotypes for different phenotypes.

### NOS2 and COX2 are predominant in tumor regions where CD3CD8 content is lower

To further define the distribution of immunosuppressive cellular phenotypes and NOS2/COX2, the ratio of immunosuppressive markers (i.e., T_reg_, Mac-PDL1, etc.) with tumor-COX2^+^ was analyzed and plotted against T_eff_ (CD3CD8PD1^-^). The highly significant linear correlation between tumor-COX2+ and tumor-NOS2+ (Fig.1C) allows either to used to evaluate their association with different cellular phenotypes. Due to the antagonistic relationship concerning survival between NOS2 or COX2 with T_eff_ cell number, a ratio of each marker to tumor-COX2^+^ was calculated and plotted against the percentage of T_eff_ cells. The ratio between each immune suppressive cellular phenotype and tumor-COX2^+^ by itself was insignificant (Suppl. Fig.2A). Nonetheless, when the ratio of these immune suppressive phenotypes to tumor-COX2^+^ was plotted against the T_eff_ cel percentage, a significant linear relationship was observed in which the ratio of lymphoid cells CD4 T_reg_, CD8 T_reg_, and CD8 T_ex_ (CD3CD8PD1^+^) (Fig. 2C). The correlations between tumor-PDL1 and macrophage-COX2 were only marginally significant, indicating a weak relationship (Fig. 2D). Overall, whole tumor PDL1 cellular phenotypes were not highly correlated with survival or related to other cellular phenotypes in this cohort (Fig. 2B). Thus, leukocyte-based immunosuppressive cells are favored in the presence of elevated lymphoid cells in the tumor (inflamed tumor), while NOS2 and COX2 predominates in the presence of lower overall CD3CD8 (cold, immune desert) indicting distinct roles in tumor immunosuppression between inflamed and cold regions.

### Spatial clustering of tumor-NOS2^+^ and tumor-COX2^+^ is a significant determinant of poor outcome in ER- breast cancer

To identify cellular neighborhoods that correlate with outcome, a spatial uniform manifold approximation and projection (S-UMAP) analysis of all single cell neighborhoods was conducted (58). The S-UMAP was applied to all single cell neighborhood density profiles constructed of 9 phenotypes (CD4^+^FOXP3^+^ CD8^+^FOXP3^+^, CD8^+^CD3^+^PD1^+^, tumor NOS2^+^, tumor COX2^+^, tumor PDL1^+^, CD68^+^PDL1^+^, CD68^+^COX2^+^, and others) to identify clusters of similar neighborhoods. The differential cluster distribution (the ratio of cluster populations by outcome) revealed that relative to survivors, deceased patients had a greater percentage of cellular neighborhoods with elevated levels of tumor-NOS2^+^, tumor-COX2^+^, CD4^+^FOXP3^+^, CD8^+^FOXP3^+^, and CD68^+^PDL1^+^ (Fig 2E, F). Comparison of distances between individual factors show that tumor-COX2^+^, and tumor-NOS2^+^had a gradient effect where the correlation with inhibitory cell types in deceased patients deceased rapidly with distance (Fig 2G). In the cellular neighborhoods of deceased patients, the tumor-COX2^+^ and tumor-NOS2^+^ densities decreased rapidly over 100 µm (Fig 2G). This suggests that clustering of tumor-NOS2^+^ and tumor-COX2^+^ in spatially distinct regions is an important determinate of poor outcome. In contrast, T_reg_ and T_ex_ (CD68^+^PDL1^+^) impact on survival was not dependent on distance (Fig 2G). Furthermore, exhausted T-cells CD8^+^CD3^+^PD1^+^and tumor-PDL1 were marginally predictive of good outcome which could be attributed to success of extensive CD8 tumor infiltration and widespread IFNγ (Fig 2G). This unbiased analysis identifies the cellular phenotypes that impact survival, implicating the presence of distinct neighborhoods in deceased patients with specific configuration and location of these specific cellular immunosuppressive phenotypes. Thus, S-UMAP analysis validates our contention that tumor-NOS2^+^ and tumor-COX2^+^ in spatially distinct cellular regions is an important determinant of poor outcome in patients with ER- breast cancer.

### Immune suppression mechanism occurs in specific regions of the TME

The correlation between T_eff_ density and immunosuppressive phenotype suggests that the mechanism of immunosuppressive factors has regional spatial orientation connotations. Previous reports show that tumor-excluded (cold) *v**s.*** infiltrating (hot) CD8 cells have an impact on survival (56). The tumor has numerous spatially distinct regions, spanning from lymphoid aggregates to tumor cores; therefore, the regional distribution of immune-tumor interactions was evaluated. To this end, five distinct regions with specifically defined tumor anatomical features were annotated, including lymphoid aggregates and small and large tumor nests. The tumor regions were subdivided into edges of a large tumor’s NOS2+/- region at the tumor/stromal interface and the interior (core) (**Fig. 3A-C**). The smaller tumor fragments were defined tumor clusters > 0.02 mm^2^ and in most cases less than 3 cells thick in one dimension. In contrast, the larger tumor nest > 0.1 mm^2^ was used to designate the tumor core or edges. Spatial analysis of these regions with cellular phenotypes presented above was analyzed in these regions (**Fig. 3** **D-E**). The percentage of cell categories in each region was determined. As anticipated, lymphoid cellular phenotypes formed a gradient from lymphoid aggregates to the tumor’s interior. As expected, the lymphoid aggregates had the highest of leukocytes, including both CD3 (either CD4 or 8) and CD68 cells, with far less tumor cells (**Fig. 3D-G**). In contrast, the tumor core had considerably lower leukocytes (<3%) and more tumor cells. One feature is that alive patients had 4 times more T_eff_ in the interior of the tumor than deceased, indicating more infiltration, consistent with previous reports (12, 56) (**Fig. 3D**). In deceased patients, there was ∼ 1% T_eff_ cells, which corresponds to an immune desert and is consistent with CD8 cell exclusion leading to poor outcome (56). As anticipated, NOS2 and COX2 were associated with tumor regions, constituting the tumor’s predominant phenotype as compared to CD3 and CD68 (**Fig. 3** **F, G**). PDL1 was detected in both tumor cells and macrophages. CD3CD4 (Treg?) was restricted to stroma and lymphoid marginal aggregates in deceased and alive patients. As anticipated, the phenotypes of these immune suppression markers were in distinct regions.

**Figure 3.**
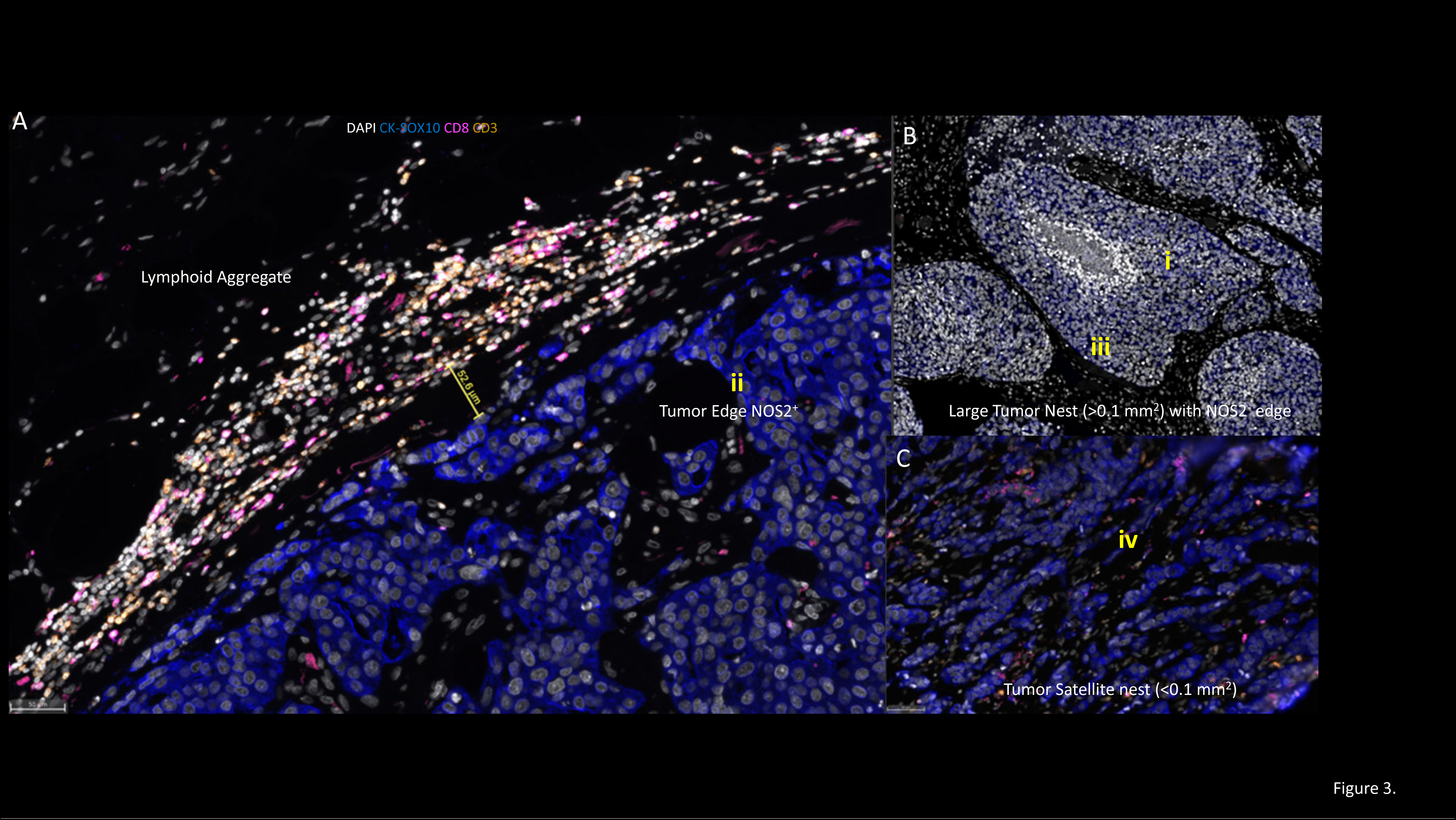
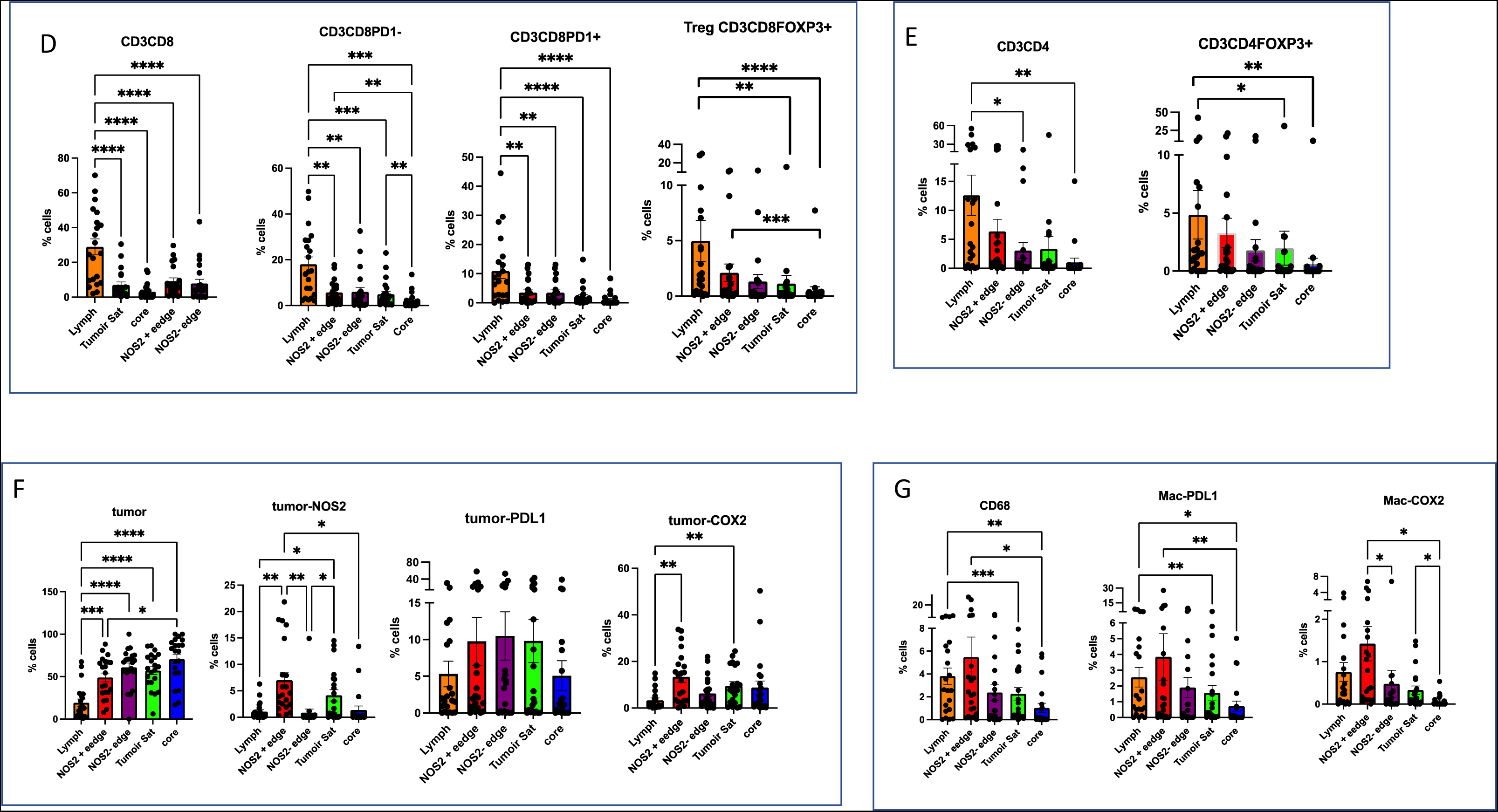
Regional placement of cellular phenotypes in the tumor. A-C) represents the 5 regions i) stromal/marginal lymphoid aggregates; Large Tumor nests (>0.1 mm^2^) with ii) NOS2^+^ tumor edges or iii) NOS2^-^ tumor edges, and iv) tumor interior (core) and tumor satellites (< 0.01mm^2^). The % cells in each of the 5 regions for cellular phenotypes D) CD8 E) CD4 F) tumor G) macrophage ((*p <0.5, **p <0.01, *** p<0.001, Ordinary one-way ANOVA).

### NOS2/COX2 drives Teff exclusion from the tumor

As shown above, the large and even small tumor nests exclude CD3CD8 cells from the core, resulting in the development of immune deserts. A previous report suggested that in TNBC, fibrosis was the primary driver of this processes, restricting mobility of CD8, while others suggest an unknown factor (56). Examining the edge of the larger tumors, NOS2 and COX2 were expressed at the tumor edge and less in the core, suggesting an essential role in excluding T_eff_. In a comparison of different regions of the different tumors, there appear to be 3 classes of CD8 T-cells exclusion from the tumor nest forming immune desert: Type I: restricted inflamed with NOS2^+^ edge and COX2^hi^; Type II NOS2^-^ with COX2^hi^ and Type III with NOS2^-^, sporadic COX2 with few proximal stromal lymphoid cells <500 µm (<1%) (figure 4A). These observations point to COX2 as having a significant role in the final phases and maintenance of the tumor’s immune desert and exclusion of T_eff_ cells.

**Figure 4.**
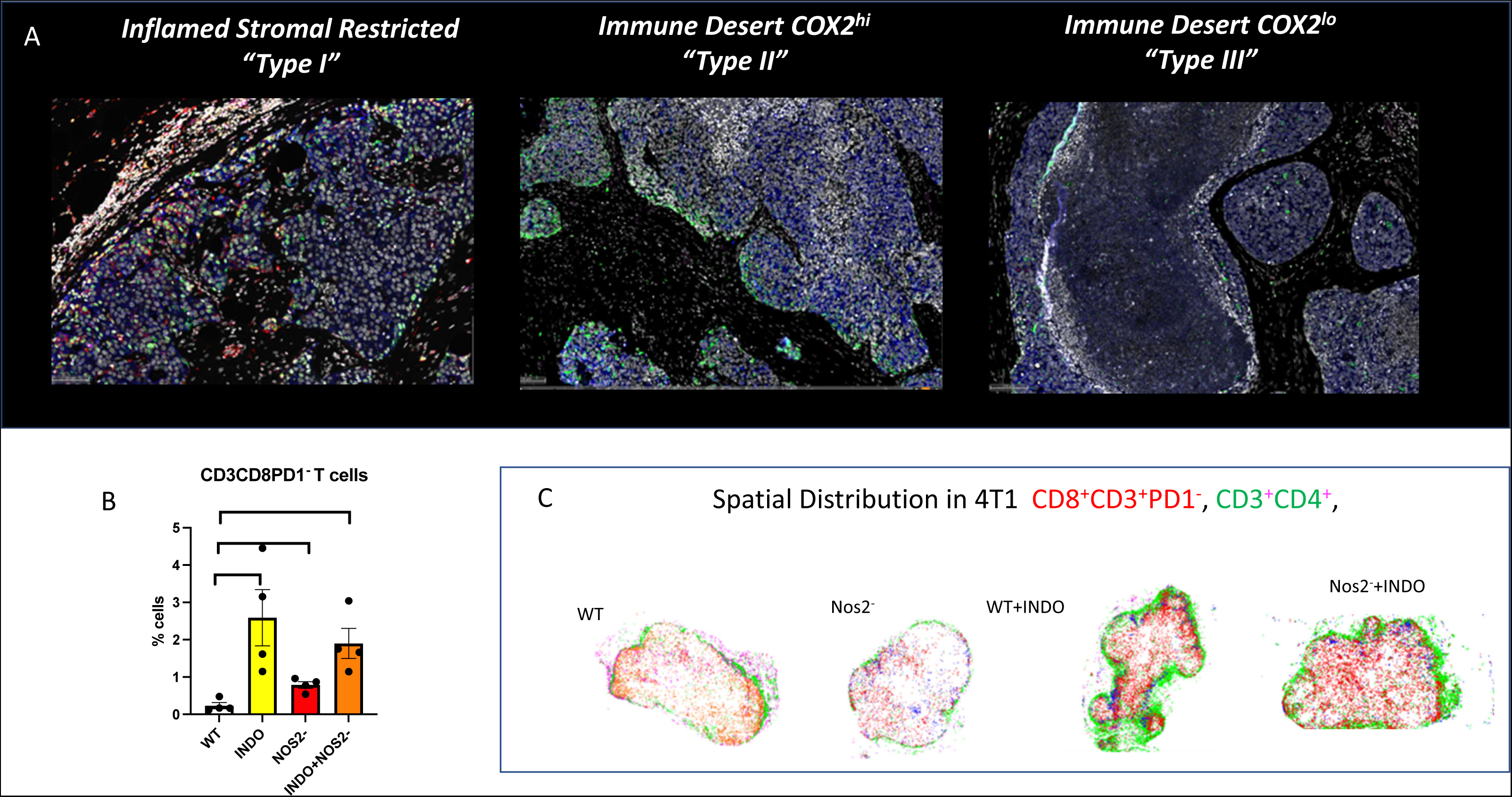
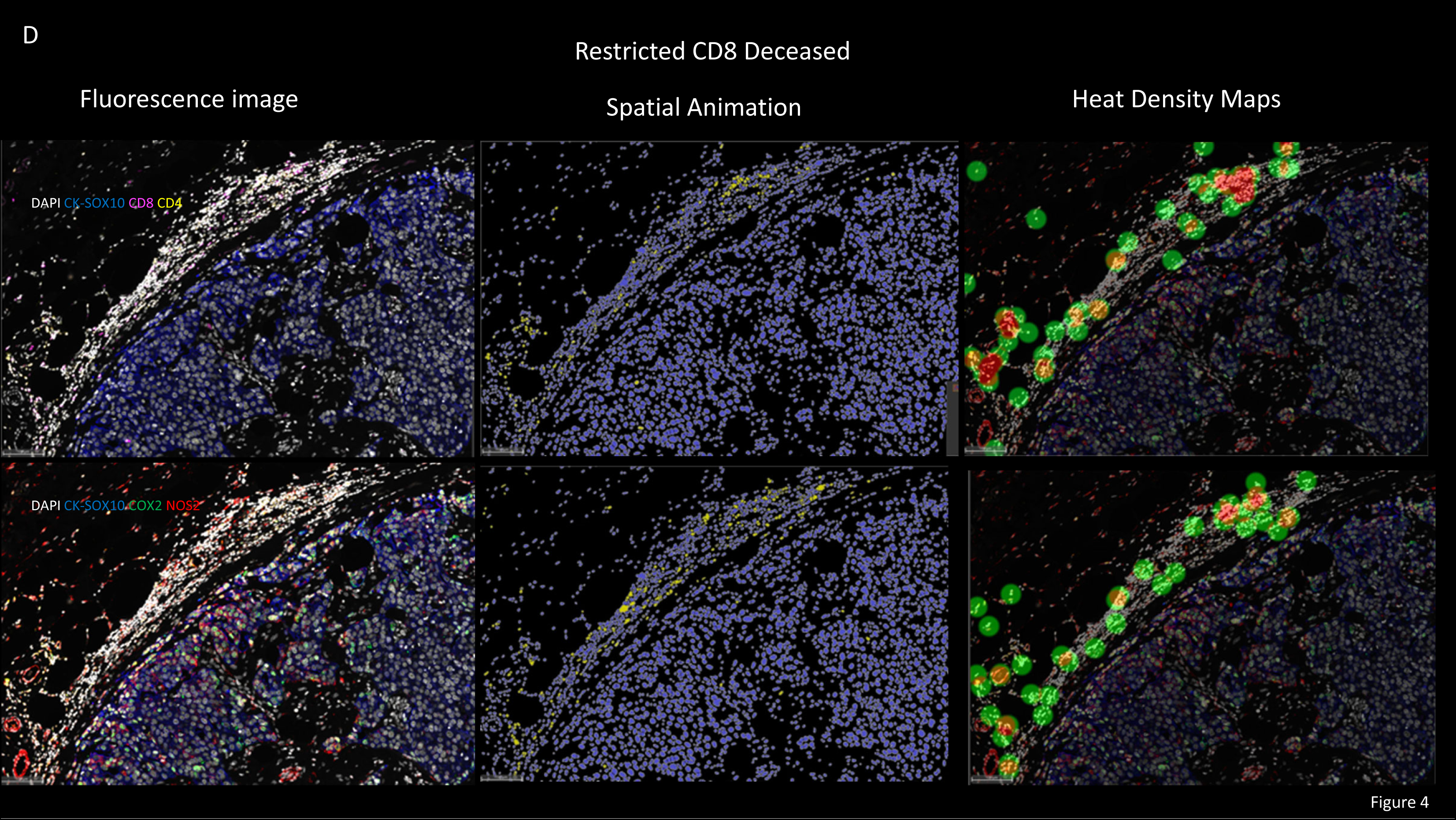
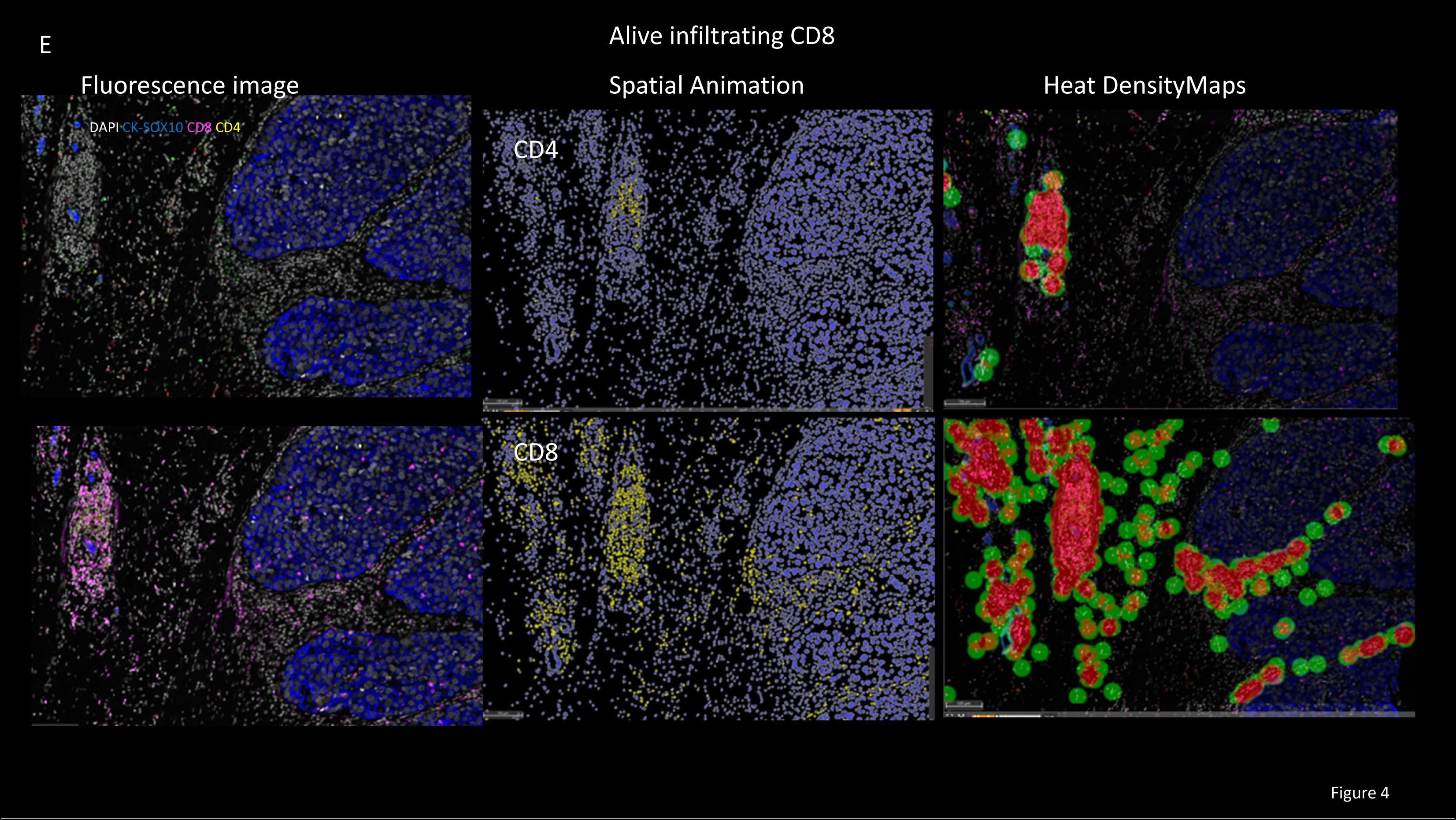
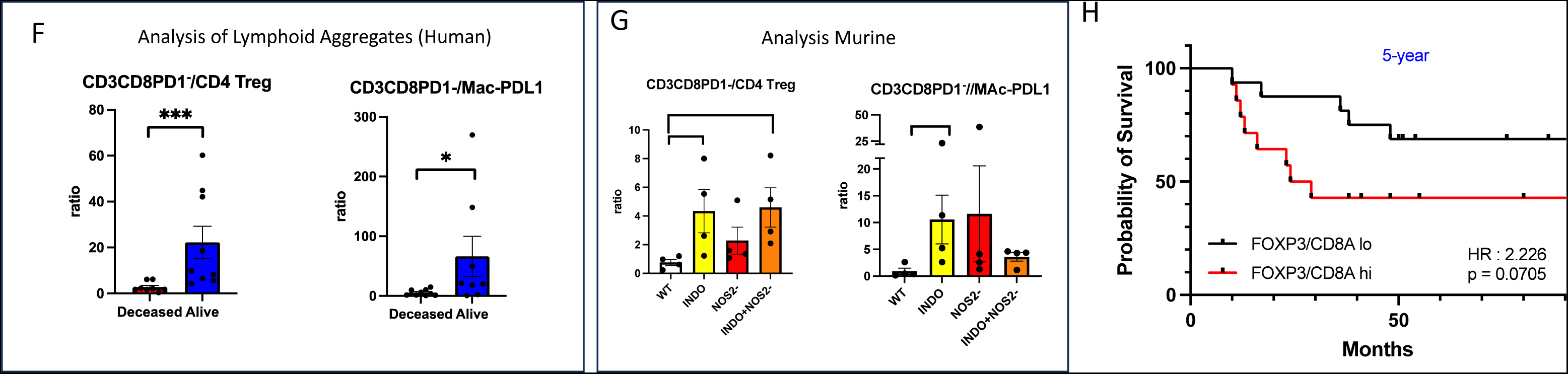
Analysis of T cell exclusion in human ER- breast cancer and 4T1 murine breast cancer. A) represents 3 types of CD8 tumor exclusion. B) is the analysis of the impact of NOS2- and indomethacin on infiltration of CD3CD8PD1- cells in 4T1 model. C) spatial distribution maps of CD3CD8PD1- (red) and CD3CD4 (green) upon treatment in 4T1 murine model. Comparison of CD4 and CD8 distribution with fluorescence imaging, annotation, and heat maps in human patients D) deceased and E) Alive. Ratio analysis of lymphoid aggregates of Teff with Treg and immune suppressive macrophage in F) human ER- breast cancer G) 4T1 tumors. H) Comparison of Kaplan Meir plot of FOXP3A/CD8A high vs FOXP3/CD8A low ratios, cohort (GSE37751) divided at median of gene expression ratio.

To investigate the role of NOS2 and COX2 in T_eff_ exclusion, a TNBC murine model was used. Our previous study shows that in the murine TNBC 4T1 model in untreated control, CD8 lymphoid cells were restricted in the margins similar to the immune desert in the human deceased patient (**Fig. 3A** **& B**) (24). While there was less difference between the control and NOS2- animals, inhibition of COX2 with indomethacin (INDO) resulted in extensive infiltration of Teff and a dramatic increase in IFNγ and CD8 cells (24). As depicted in Figure 4B, spatial analysis reveals the infiltration of T_eff_ (CD3CD8PD1^-^) cells into the core of the tumor, resulting primarily from COX2 inhibition less from NOS2. In contrast, CD4 cells remained on the outside despite T_eff_ infiltration(24) (Fig. 4C). This evidence suggests that while NOS2 and COX2 control the migration of CD8, CD4 mobility is unaffected. These results show that COX2 has a significant role in excluding CD8 cells from the tumor’s interior and that targeting it can convert cold tumors to inflamed tumors with infiltrating CD8 T_eff_, resulting in a more effective response.

The differential movement of CD8 and CD4 observed in the murine model was examined in humans, comparing deceased and alive patients. Regions at the tumor edge proximal to a lymphoid patch were examined in type I immune desert (inflamed but stromal restricted T_eff_) correlating with deceased patients compared with alive patient infiltrating T_eff_. As seen in Figure 4D, the deceased patient with restricted CD8 and CD4 were predominately confined to the lymphoid aggregate. However, in the alive patient the CD8 readily migrated, infiltrating the tumor core, yet like in the murine model, the CD4 cells remained in the lymphoid aggregate. Taken together, from murine and human samples clearly show that COX2 controls CD8 mobility while CD4 cells remain near the lymphoid aggregate (**Fig. 4E**).

COX2/PGE2 have been shown to differentially impact various types of T cells, favoring more immune suppressive phenotypes. When the lymphoid aggregate was analyzed for different immune cellular phenotypes, there was a difference between deceased and alive patients. Analysis of T_eff_ ratio with different immune markers comparing survival showed only the ratio of T_eff_/CD4 T_reg_ and T_eff_/Mac-PDL1 had a significant effect on survival (**Fig. 4F**). This finding was corroborated in the 4T1 model where addition of indomethacin and INDO/NOS2- inhibition resulted in higher T_eff_/CD4 T_reg_ or T_eff_/Mac-PDL1 ratios (**Fig. 4G**). To further extend this observation, CD8a and FOXP3 showed increased HR, validating these ratio relationships (**Fig. 4H**). The above data taken together clearly show that COX2 and NOS2 have a significant role in controlling the infiltration and mobility of CD8 as well as the immune polarization within the tumor lymphoid aggregates.

### NOS2 and COX2 are spatially distinct from Treg and PDL1

Figures 2 and 3 show that NOS2 and COX2 are distinctly localized. Therefore, a comparison of different types of whole tumor configurations was undertaken. Whole tumor phenotypes were established based on the distribution of CD3CD8PD1-, tumor-NOS2, and tumor-COX2 (Suppl. Fig 3A). Correlation analysis was used to determine the threshold based on the % of cells in the whole tumor as high (+) and low (-), which provided the development of general whole tumor phenotypes (Suppl. Fig. 3A-C). As seen from a scatter plot of NOS2 or COX2 versus T_eff_, there are 3 distinct major phenotypes (sup. Fig 3B). In the surviving patients, the major phenotype was T_eff_^+^ CD8^+^ (CD3CD8PD1- (9/10)) which upon closer examination shows infiltrating CD8 cells (Fig.4A). Conversely, deceased patients had more varied distribution ranging from CD8^+^NOS2^+^COX2^+^ (restricted CD8 inflamed type I immune desert (fig 4A)) to CD8^-^NOS2^-^ COX2^lo^ (Type III immune desert (Fig. 4A). The majority of deceased patients exhibited a predominant CD8^-^NOS2^-^COX2^+^ (type II immune desert, Fig. 4A) phenotype. The difference between these phenotypes is that CD8 cells were restricted from the tumor in deceased patients while the tumors were fully infiltrated in survivors.

Next, density heat maps of tumor-PDL1, T_reg_ FOXP3, tumor-NOS2^+^, and tumor-COX2^+^ compared with CD3CD8 were done to identify the topography of each cellular neighborhood (Fig. 5). Restricted tumors classified as CD8+NOS2+COX2+ had two distinct regions containing both high (restricted inflamed) and devoid CD3CD8 regions (immune desert) (Fig. 5A). Figure 5A shows that in restricted CD8 regions, lymphoid aggregates also had T_reg_ and PDL1. In contrast, regions with low or absent CD3CD8 cells contain more tumor-NOS2^+^and tumor-COX2^+^, consistent with figures 2 and 3, supporting the existence of regional differences. One deceased patient with inflamed but restricted CD8+ was devoid of either NOS2 or COX2. However, extraordinarily high levels of T_reg_, PDL1, and PD1 were observed. Thus, confirming the notion that T_reg_, PDL1, and PD1 are spatially and even temporally associated with T_eff_ cells while orthogonal to NOS2 and COX2 (Fig 5B). This clearly supports the linear analysis in Fig. 2C. Figures 5C and D represent Low CD3CD8 (immune deserts) with low NOS2 but higher COX2. In contrast, alive patients have low levels of NOS2 and COX2, and high levels of infiltrating CD3CD8PD1- yet there were low levels of T_reg_, implicating that in the absence of NOS2/COX2, T_reg_ is a predominate factor in immune suppression.

**Figure 5.**
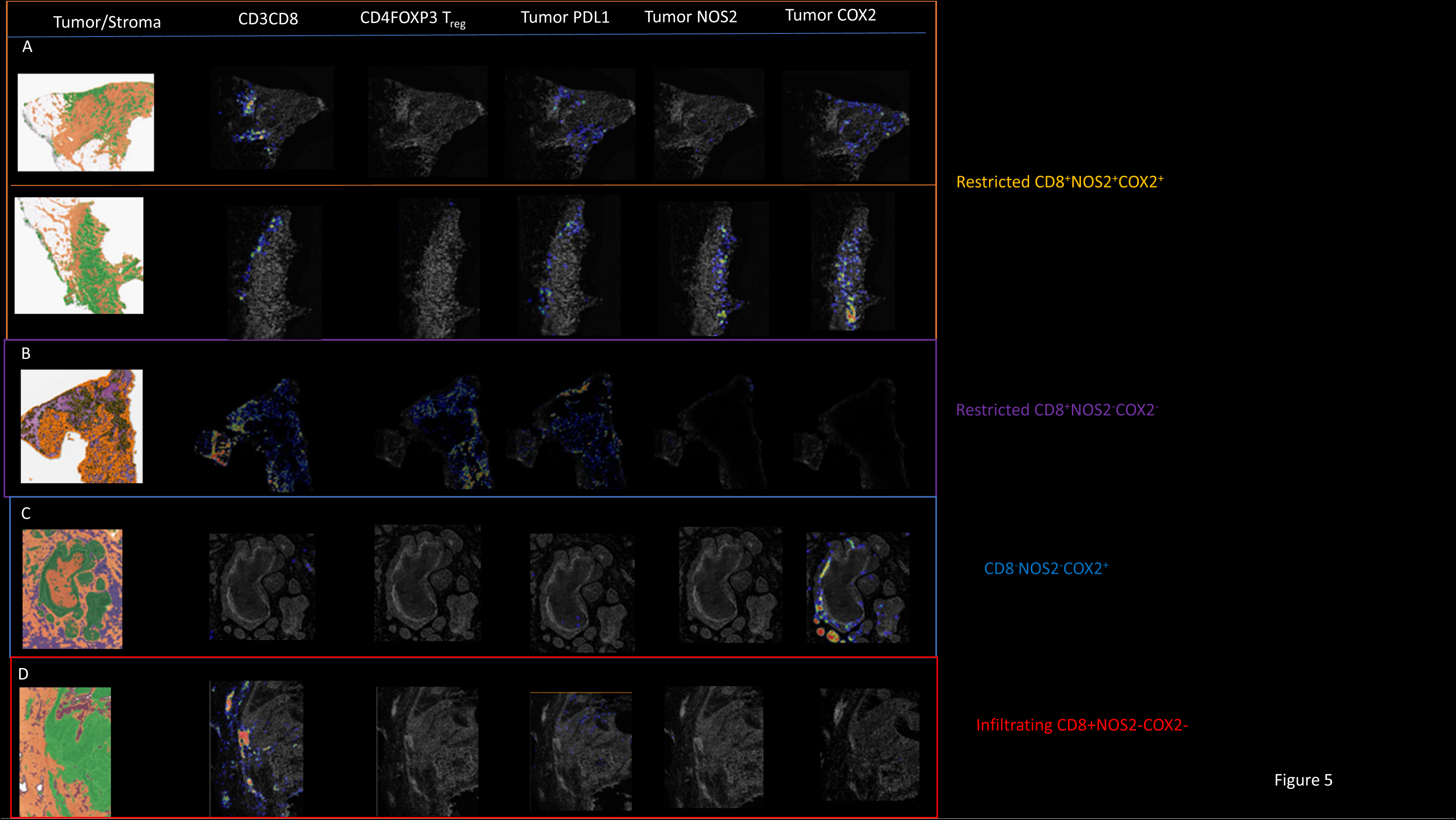
The density heat map distribution comparisons. The density heat map distribution comparing CD3CD8, CD4FOXP3^-^Treg, tumor-PDL1, tumor- NOS2 and tumor-COX2 in whole tumor classification of CD8NOS2 and COX2. A) restricted CD8^+^NOS2^+^COX2^+^ B) restricted CD8^+^NOS2^-^COX2^-^ C) CD8^-^NOS2^-^COX2^+^ D) infiltrating CD8^+^NOS2^-^COX2^-^.

In contrast, low T_reg_ with higher NOS2 and COX2 leads to immune suppression and poor outcomes. In addition, there were low levels of PDL1 and high levels of IFNγ. Since PDL1 can be induced by IFNγ expression, this indicates the efficacy of tumor eradication. These two distinct profiles of deceased and living patients demonstrate that NOS2 and COX2 play a significant role in determining survival and is involved from the transition from inflamed to the immune desert.

### IDO1 and B7H4 have distinct spatial orientation

Previous studies show that the immunosuppressive factors IDO1 and B7H4 were in distinct regions (56). Whereas IDO1 was largely associated with the lymphoid regions, the B7H4 was associated with the tumor core (56, 59). Here, spatial analysis using multiplexing fluorescence for IDO1 and B7H4 were examined and compared with other immunosuppressive markers. When IDO1 and B7H4 were evaluated in univariant survival analyses, there was no significance (Fig. 6A, B). However, the ratio of NOS2s to IDO1 and B7H4 was significant relative to T_eff_ and IFNγ (Fig. 6A, B). Since IDO1 and B7H4 can be induced by IFNγ (60), this may reflect T_eff_/IFNγ activity. Previous studies indicate that a high IDO1 level is a favorable prognostic indicator for ER^+^ but not significant for ER- breast cancer (61). These findings suggest that NOS2/COX2 and IDO1 have a contrary relationship in terms of survival. IDO1 to NOS2 versus CD8CD3 demonstrates, once more, that the IDO1/NOS2 ratio is associated with elevated CD3CD8 and, more importantly, CD8CD3PD1- cytolytic T cells.

**Figure 6.**
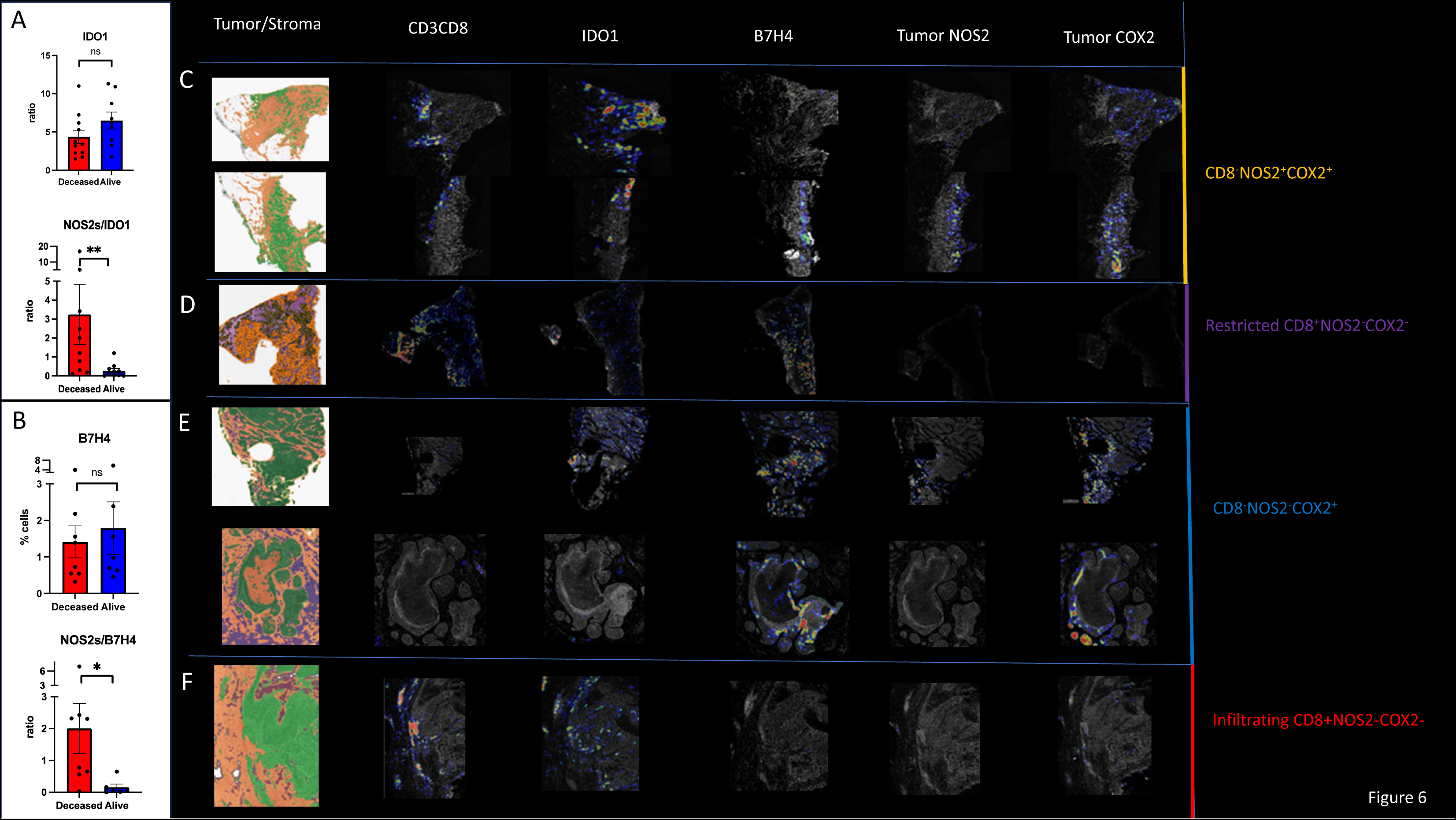
The survival and spatial analysis of IDO and B7H4. A) Compares survival as univariant the ratio of NOS2s with IDO1. B) Compares survival as univariant the ratio of NOS2s and B7H4. (*p <0.5, **p <0.01, *** p<0.001, Mann-Whitney test, one-tail). C) Density heat maps of CD3CD8, IDO1, B7H4 comparing to tumor-NOS2 and tumor-COX2. D-F) Density heat maps of CD3CD8, IDO1, B7H4, tumor-NOS2, and tumor-COX2 in restricted CD8^+^NOS2^-^COX2^-^, CD8^-^NOS2^-^COX2^+^, and Infiltrating CD8^+^NOS2^-^COX2^-^ regions.

Next, the localization of IDO1 and B7H4 was compared to CD3CD8, tumor-NOS2 and tumor-COX2. Using density heat maps, it is confirmed that higher IDO1 concentrations are proximal to lymphoid-restricted regions, indicating association with the lymphoid aggregates (Fig 6C-F). However, B7H4 was associated with immune desert and the core of the tumor. Comparing these to tumor-NOS2 and tumor-COX2, IDO1 was more closely associated with edge and lymphoid aggregates proximal to NOS2 in deceased and widely distributed in alive patients. In contrast, the immune desert contained both B7H4 and COX2. Thus, IDO1 is associated with inflamed regions containing CD3CD8 cells proximal to NOS2 in deceased patients, but the immune desert is predominately COX2 and B7H4. A landscape of different immunosuppressive cellular phenotypes shows those that are predictive of survival occupying unique regions of the tumor microenvironment.

## Discussion

Poor outcomes for cancer patients are determined by processes such as metastasis, chemoresistance, and immunosuppression. The immunosuppressive tumor microenvironment is a significant obstacle to successful immunotherapy, thus interventions capable of reducing immunosuppression are promising. Our study shows that that NOS2 and COX2 have powerful immunosuppressive properties that participate in progressive depression of the immune antitumor response. IFNγ from T_eff_ is crucial in inducing NOS2/COX2 expression in TNBC (12), and the resulting NO/PGE2 production inhibits T_eff_ function. This dichotomy represents a classic negative feedback mechanism, where IFNγ from activated T_eff_ increases tumor-NOS2 and COX2, dampening the Th1 antitumor response. The specific spatial orientation of T_eff_/NOS2/COX2 is a *critical determinant of outcome and shapes the immunosuppressive response*. Patients with positive outcomes have low, sporadic, NOS2 and COX2 expression, with T_eff_ penetrating the tumor core. In contrast, patients with stromal or marginally restricted T_eff_ can have elevated NOS2 and/or COX2, which, when regionally clustered, forms a leukocyte-depleted zone that does not permit T_eff_ penetration. Thus, the spatial relationship between NOS2/COX2 expression and T_eff_ cells shapes the immune environment of the TME.

There are 3 distinct types of CD8 T-cell exclusion from the tumor core with respect to T_eff_/NOS2/COX2 that correlate with poor outcomes: **Type I**, *restricted inflamed*, with stromal restricted CD8 T-cell inflamed regions with elevated tumor NOS2 and COX2; **Type II**, *developing immune desert* with a lower percentage of stromally restricted CD8 T-cells where there is tumor-COX2^+^ without tumor-NOS2; and **Type III**, *indicating mature immune desert* with <1% CD8 cells (<500 µm (<1%) with tumor-NOS2^-^, and sporadic COX2 with few stromal lymphoid cells with elevated B7H4. Another feature that distinguishes Type I and II from III is that in Type I and II there are proximal lymphoid aggregates to the tumor, while in type III there is none. Although these are distinct regional phenotypes, it is not uncommon to observe all of them in the same tumor, suggesting that these are linked through a progressive process from the inflammatory entry of tumor infiltrating lymphocytes (TIL) inducing NO/PGE2 production, leading to mature immune deserts.

CD8 exclusion from the tumor prevents the cell-to-cell contact necessary for T_eff_ release of cytolytic effector molecules into tumor cells. This specific cell interaction targets tumor/virally infected cells, limiting damage to normal tissue in contrast to the non-specific killing of rapidly growing cells that occurs with conventional therapies (62–65). Thus, interventions that enhance T cell entry into the tumor and their tumoricidal activity will be therapeutically beneficial. In a previous study in TNBC, restriction of CD8 was related to poor outcomes, consistent with other reports in various cancers; therefore, higher TIL can be restricted or infiltrated (66–69). Furthermore, the authors show that there were three different types of CD8 exclusion: stromally restricted, marginally restricted, and complete immune desert (<1% CD8). However, an understanding of the distinct spatial distributions is required to clearly define the mechanisms operating in the immune/tumor interaction. The definition of the three types of CD8 exclusion regarding NOS2 and COX2 expression shown here demonstrates that tumor exclusion of T_eff_ is controlled by NOS2 and COX2.

While NOS2 and COX2 impact the whole tumor landscape, they also influence specific tumor regions, increasing the TME heterogeneity. The 5 regions: large tumor nests with or without NOS2 expression, lymphoid aggregate, tumor edge, tumor core, and tumor satellites, are impacted differently by the spatial configuration and distribution of T_eff_/NOS2/COX2. The regional restriction maintaining distance between lymphoid cells appears to be an essential process to protect the tumor and reduce the TIL. More importantly, each region has a distinct immunosuppressive mechanism. For example, the core of larger tumors is predominated by COX2 and B7H4, while the inflamed edge is NOS2^+^/COX2^+^. In the lymphoid aggregates, immune cell-based immunosuppression predominates, such as T_reg_ and Mac-PDL1. The specific regions of immunosuppression relative to T_eff_/NOS2/COX2 spatial configuration provide insight into the role of NOS2 and COX2 in general immunosuppression.

Since the spatial defined immunosuppressive niches share common mechanisms, they indicate that complimentary use of immune activating therapy may have synergistic effects targeting different regions. In the Type I configuration, the increased restricted lymphoid cells reside primarily in the lymphoid aggregates. The immune polarization of this region can impact survival. The survival comparisons of T_eff_ with other markers, specifically in these regions, showed two phenotypes predicting poor outcomes: CD4 T_reg_ and Mac-PDL1. The 4T1 model also shows this trend, with NOS2 or COX2 inhibition increasing the ratios of T_eff_ to CD4-T_reg_ and T_eff_ to Mac-PDL1 associated with improved survival, demonstrating that NOS2 and COX2 determine the nature of the lymphoid aggregates of TIL.

In various types of cancer, high T_reg_ cells and a low ratio of CD8^+^ T cells to T_reg_ cells in the TME are associated with unfavorable prognosis (70–72). The T_reg_ phenotype is induced by TGFβ, IFNγ, and IL4 (73–75), driving IDO1 and kynurenine production further increasing T_reg_ cells. T_reg_ produce numerous immunosuppressive cytokines, including IL10 and TGFβ. Both PGE2 and NO can increase T_reg_ factors such as IL10. T_reg_ cells suppress T_eff_ through direct PGE2 stimulation of IL10. With NOS2 and COX2 driving cytokine release and activation of IL10 and TGFβ, these systems complement each other regionally to maintain immune suppression.

While tumor-PDL1 expression can be located at the tumor-stromal interface, the Mac-PDL1 is an immunosuppressive tumor-associated macrophage (TAM) that resides primarily in the lymphoid aggregates. Mac-PDL1 expression occurs through exposure to IFNγ, TNF, and IL6 as part of a negative feedback loop (76, 77). TNF-ɑ and IL-6 activate the NF-kB and STAT3 signaling pathway to regulate PD-L1 expression on M2 TAM (76–79). Increased PDL1 expression on macrophages is involved in developing the TAM phenotype (77, 80). Mac-PDL1 has been shown to inhibit T_eff_ in collaboration with IDO1 or granulin (81–85). The IDO1 density heat maps show a strong association with high CD3CD8 regions, suggesting that an increase in this M2 macrophage phenotype relative to T_eff_ indicates poor CD8 health. Thus, the increase in different cytokines and proximity to IDO1 suggest that poor outcomes is due in part to the induction of the M2 immunosuppressive phenotype that polarizes the lymphoid aggregate.

***Type I restricted-inflamed tumors*** show increased tumor-NOS2 and -COX2. The relative positioning of NOS2 and COX2 to T_eff_ suggests an immunosuppressive progression from inflamed foci to an immune desert devoid of lymphocytes. The NOS2^+^ regions with complimentary COX2^+^ can increase prooncogenic pathways and lead to metastasis. NOS2 and NO have been shown to increase metastasis. The higher levels of NOS2 lead to an increase in oncogenic pathways and immune modulatory proteins such as IL1, IL6, IL8, and TNFα (86). Regions of elevated NOS2 and COX2 would be expected to increase cancer cell stemness, thus leading to therapy resistance while also causing increased activation of TGFβ and IL-10. The diffusion of these immunosuppressive factors would counter Th1 inflammation, favoring more immunosuppression. Thus, the Type I configuration is highly inflamed with distinct areas that fuel immunosuppression through complementary mechanisms. Therefore, the interaction of lymphoid aggregates and the tumor edge leads to immune suppression and increases chemoresistance and metastasis, the hallmark of poor outcomes.

***Type 2 and Type 3 immune deserts*** represent regions with decreased TIL having a CD8-NOS2-COX2± phenotype. These are immunologically cold tumor regions, with Type 2 having lymphoid aggregates proximal (< 0.5 mm) while Type 3 does not. Type 2 shows a gap between many of the tumor nests and leukocytes that is 50 -100 µm. High COX2 rather than NOS2 suggests the importance of PGE2 from the tumor in maintaining the exclusion of CD8 cells. The lack of TIL aggregates in type 3 is associated with lower COX2 levels together with B7H4 expression. Thus, these regions of the immune desert may be predisposed to therapy resistance. The 4T1 model produces these Type 2/3 immune deserts and has features that resemble human tumors. The addition of radiation simply augments the marginal immunosuppressed populations, despite the cell injury. However, NOS2 and COX2 inhibition activates T_eff_, increasing granzyme B. The enrichment of Type 2/3 immune phenotypes through damage due to treatment or tumor stress such as hypoxias and ischemia/reperfusion can recruit lymphocytes that produce IFNγ and cytokines. However, they also induce a robust NOS2/COX2 immunosuppression response, thus leading to poor clinical outcome. Several studies show that more advanced tumors have poorer responses and reduced TIL penetration into the tumor core (87–90). This observation would reflect a fortified tumor barrier where preferential activation of NOS2/COX2 dramatically limits antitumor immune response.

In more advanced tumors such as chemoresistant breast cancer, the combination of tumor cell chemoresistance and immunosuppression results in poor outcomes. A recent study of chemoresistant TNBC patients treated with a pan–NOS inhibitor low dose aspirin showed remarkable positive outcomes with >88% response in locally advanced tumors (91). The administration of the NOS inhibitor resulted in changes in the TME where there was some increase in CD8 together with M1 macrophages, neutrophils (N1), and B cells (91). Remarkably similar changes were observed in the 4T1 model, where NOS2-/INDO led to increased CD8 cells, N1, and B cells, resulting in cures (24). In the murine TNBC 4T1 model, NSAID alone only targeted the T cells, providing a delay but not a cure. This remarkable similarity between the clinical trial and animal model suggests the importance of targeting NOS2 and COX2 to relieve immune suppression. Crosstalk between NOS and COX2 leads to chemoresistance and cancer stem cells while mediating immunosuppression.

## Conclusion

NOS2 and COX2 tumor expression shape the immune landscape and mediate the transition from inflamed regions to immune deserts. While NOS2 and COX2 have numerous direct suppressive mechanisms, they also enhance other immunosuppressive mechanisms. It is important to appreciate the distinct regional mechanisms of immune suppression resulting from the differential expression of NOS2 and COX2. The formation of an immune desert and the subsequent resistance to treatment linked to NOS2/COX2 induction in the tumor indicate that novel treatments targeting these proteins may have value for more advanced therapy-resistant tumors. NOS2 and COX2 augment multiple immunosuppressive pathways and increase the severity of immunosuppression (Fig.7). Taken together, these data show that in ER- breast cancer, NOS2 and COX2 have a prominent role in immunosuppression making it treatment resistant. Thus, NOS2 and COX2 targeting may increase the efficacy of a wide range of treatments.

**Figure 7.**
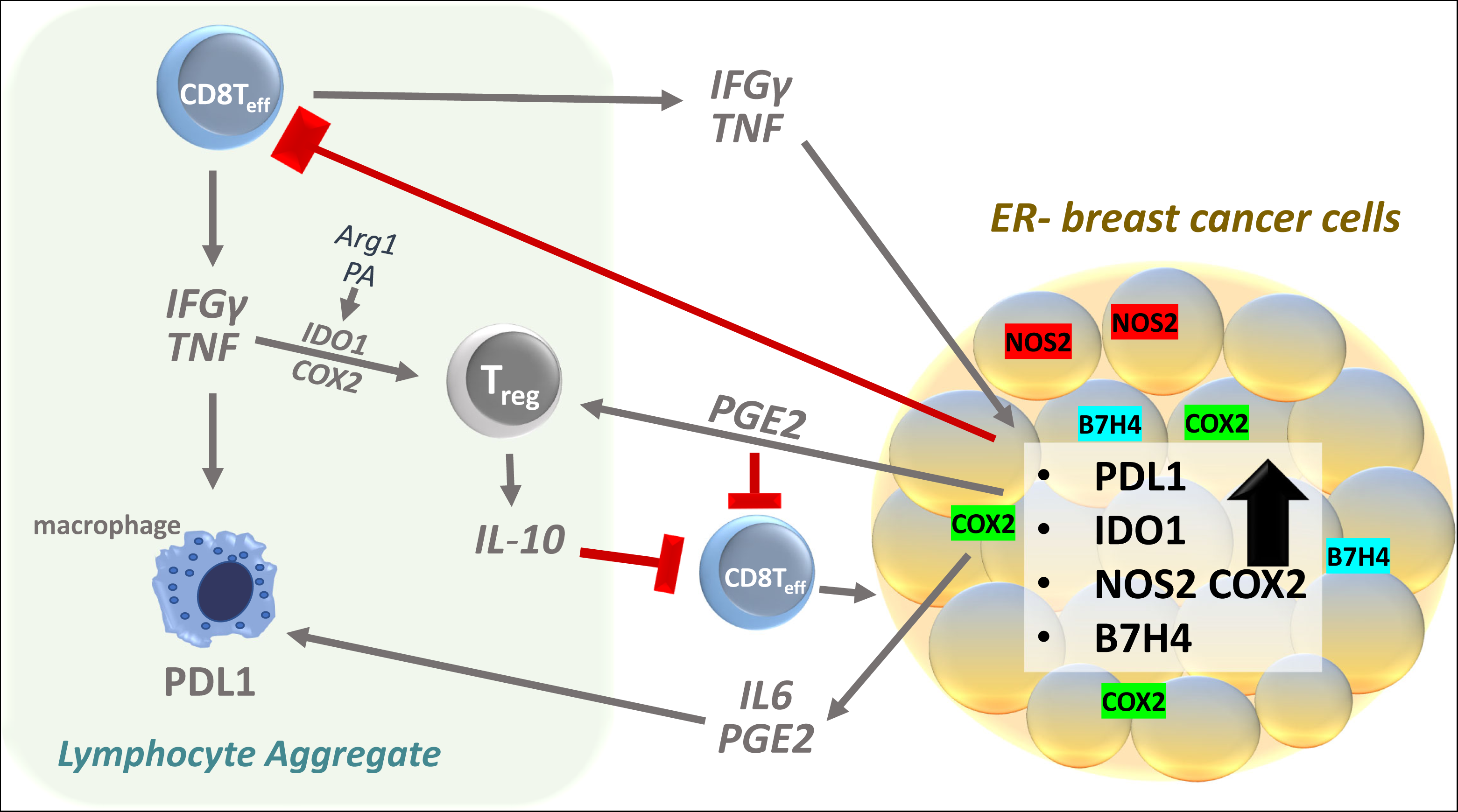
Graphic analysis of the impact of T cell interaction with NOS2 and COX2 in the tumor microenvironment and mechanism that cooperate to lead to immune suppression.

Determining the immunosuppressive phenotypes of distinct regions of the tumor landscape provides insight into the multiple local mechanisms affecting tumor survival. The introduction of checkpoint inhibitors has provided a new approach to the treatment of various malignancies. In breast cancer, however, checkpoint inhibitors have been significantly less effective (55, 92, 93). Immunosuppressive factors that reduce the efficacy of many cancer treatment strategies present a major obstacle to successful therapy. According to the data presented herein, restriction and even elimination of CD8 effector cells by NOS2 and COX2 suggests that NO and PGE2 play an essential role in immunosuppression and agents that control their activity offer an important clinical tool in controlling the immune response.

## Materials and Methods

### Tissue Collection and Immunohistochemical Analysis of Patient Tumor Sections

Tumor specimens (n = 21) were obtained from breast cancer patients recruited at the University of Maryland (UMD) Medical Center, the Baltimore Veterans Affairs Medical Center, Union Memorial Hospital, Mercy Medical Center, and the Sinai Hospital in Baltimore between 1993 and 2003. Informed consent was obtained from all patients. The collection of tumor specimens, survey data, and clinical and pathological information (UMD protocol no. 0298229) was reviewed and approved by the UMD Institutional Review Board (IRB) for the participating institutions. The research was also reviewed and approved by the NIH Office of Human Subjects Research (OHSR no. 2248). Breast tumor NOS2 and COX2 expression was analyzed previously by IHC using 1:250 diluted NOS2 antibody and 1:50 diluted COX2 antibody (no. 610328 and 610204, respectively, BD Biosciences, San Diego, CA,) and scored by a pathologist (49, 50). For NOS2 staining, a combination score of intensity and distribution were used to categorize the immunohistochemical NOS2 stains where intensity received a score of 0-3 if the staining was negative, weak, moderate, or strong. The NOS2 distribution received scores of 0-4 for distributions <10%, 10-30%, >30-50%, >50-80% and >80% positive cells (49). For COX2 staining, scores of negative to weak (1–2) or moderate to strong (3–4) were categorized as low or high, respectively (50). Herein, NOS2 and COX2 expressions were also analyzed by fluorescent staining performed on the Leica Biosystems (Wetzlar, Germany) Bond RX Autostainer XL ST5010 using the Bond Polymer Refine Kit (Leica Biosystems DS9800), with omission of the Post Primary Block reagent, DAB and Hematoxylin. After antigen retrieval with EDTA (Bond Epitope Retrieval 2), sections were incubated for 30 min with COX2 (Cell Signaling Technology, Danvers, MA, no. 12282, 1:100), followed by the Polymer reagent and OPAL Fluorophore 520 (AKOYA, Marlborough, MA). The COX2 antibody complex was stripped by heating with Bond Epitope Retrieval 2. Sections were then incubated for 30 min with NOS2 antibody (Abcam no. ab15323, 1:50), followed by the Polymer reagent and OPAL Fluorophore 690. The NOS2 antibody complex was stripped by heating with Bond Epitope Retrieval 2 and then stained with CD8 (Abcam no. 101500, 1:100) or IFNγ (Abcam no. 231036, 1:200), followed by the Polymer reagent and OPAL Fluorophore 570. Sections were stained with DAPI and coverslipped with ProLong Gold AntiFade Reagent (Invitrogen). Images were captured using the Aperio ScanScope FL whole slide scanner (Leica). The original IHC previously reported (49, 50) and fluorescent NOS2/COX2 staining results were generally consistent. (IDO1 AND B7H4),

Formalin-fixed paraffin embedded (FFPE) tissue sectioned at 4 μm and mounted on SuperFrost Plus slides were stained with a FixVUE Immuno-8^TM^ Kit (formerly referred to as UltiMapper® kits (Ultivue Inc., Cambridge, MA), USA; CD8, NOS2, COX2, CKSOX10, and IFNγ cocktail) using the antibody conjugated DNA-barcoded multiplexed immunofluorescence (mIF) method (1). These kits include the required buffers and reagents to run the assays: antibody diluent, pre-amplification mix, amplification enzyme and buffer, fluorescent probes and corresponding buffer, and nuclear counterstain reagent. Hematoxylin and Eosin (H&E) and mIF staining was performed using the Leica Biosystems BOND RX Autostainer. Before performing the mIF staining, FFPE tissue sections were baked vertically at 60-65 °C for 30 min to remove excess paraffin prior to loading on the BOND RX. The BOND RX was used to stain the slides with the recommended FixVUE (UltiMapper) protocol. During assay setup, the reagents from the kit were prepared and loaded onto the Autostainer in Leica Titration containers. Solutions for epitope retrieval (ER2, Leica Biosystems cat# AR9640), BOND Wash (Leica Biosystems cat# AR9590), along with all other BOND RX bulk reagents were purchased from Leica). During this assay, the sample was first incubated with a mixture of all 4 antibody conjugates, next the DNA barcodes of each target were simultaneously amplified to improve the sensitivity of the assay. Fluorescent probes conjugated with complementary DNA barcodes were then added to the sample to bind and label the targets; Next, a gentle signal removal step was used to remove the fluorescent probes of the markers. The stained slides were mounted in ProLong Gold Anti-Fade Mountant (Thermo Fisher Scientific, Waltham, MA, cat# P36965) and coverslipped (Fisherbrand Cover Glass 22 x 40mm, #1.5). Digital immunofluorescence images were scanned at 20X magnification. Images were co-registered and stacked with Ultivue UltiStacker software. The digital images were then analyzed using HALO image analysis platform (94).

Previous studies indicate that NOS2 fluorescence intensity thresholds of total tumor intensity are associated with a worse prognosis. Here, the intensity levels of NOS2 and COX2 at the single-cell level were determined (see methods). The thresholds of strong, medium, and weak were determined as previous described based on average cell intensity and increasing thresholds based on standard deviation, with each value compared to significance between the deceased and alive (12, 24); (see Materials and Methods). NOS2 demonstrated that differences in survival were statistically strengthened demonstrating fourfold significance between the deceased and the alive (Fig. Sup. 1A). In contrast, the maximum difference and statical significance of COX2 did not change over the 3 thresholds. Thus, the thresholds for NOS2^s^ and NOS2^all^ were based on the analysis of NOS2^s^ and NOS2^all^, while COX2^all^ was used to represent this protein. The all designation is a sum samples of the cells that have intensity > the weak threshold limit. These findings confirm that NOS2 scoring by a pathologist is consistent with fluorescence imaging and that the elevated NOS2 and COX2 was larger associated with tumor (94).

### The Cancer Genome Atlas (TCGA) and Gene Expression Omnibus (GEO) Analyses

The breast cancer (BRCA) subset of TCGA (https://www.cancer.gov/about-nci/organization/ccg/research/structural-genomics/tcga) was accessed through the UCSC (University of California, Santa Cruz) Xena Browser (Date of access: 11/2/2022, https://xena.ucsc.edu/). GEO dataset, GSE37751 was download from the GEO portal (https://www.ncbi.nlm.nih.gov/geo/). The gene expression matrix files from both datasets were parsed and merged with their index file after download. Gene expression data, ratios between genes, and survival data were exported to GraphPad Prism (10.1.2) for survival plotting.

### Statistical Analysis

Experiments were assayed in triplicate unless otherwise stated. Student t test was test was employed to assess statistical significance using the GraphPad Prism software (version 9). Image analyses are reported as mean ± SEM and T-tests with Welch’s or Mann Whitney correction were used when appropriate to determine significance. Linear analyses and Pearson’s correlations were also conducted to determine significant correlations between protein expressions using Prism software (10.1.2). Significance is reported as *p ≤ 0.05, **p ≤ 0.01, ***p ≤ 0.001, ****p≤0.0001.

## Acknowledgements

This project was funded in whole or in part with Federal funds from the Intramural Research Program of the NIH, National Cancer Institute, CCR, CIL. This project has been funded in part with Federal funds from the Frederick National Laboratory for Cancer Research, National Institutes of Health, under contract HHSN261200800001E (ALW, WFH, DB, EFE, MP, SKA, SJL). This project was funded in part by São Paulo Research Foundation (FAPESP) grants 2018/08107-2 and 2021/14642-0 (LC, MCR), NIH R01CA238727, NIH U01CA253553, and John S Dunn Research Foundation (STCW), NCI grant no. U54 CA210181, the Breast Cancer Research Foundation (BCRF), the Moran Foundation, Causes for a Cure, philanthropic support from M. Neal and R. Neal, and the Center for Drug Repositioning and Development Program (CREDO) (JCC), Science Foundation Ireland (SFI) grant number 17/CDA/4638, and a SFI and European Regional Development Fund (ERDF) grant number 13/RC/2073 (SAG). We wish to thank the São Paulo Research Foundation (FAPESP) for student support (LC). The content of this publication does not necessarily reflect the views or policies of the Department of Health and Human Services, nor does mention of trade names, commercial products, or organizations imply endorsement by the US Government.

## Abbreviations

ARG1: Arginase 1
B7H4: a.k.a. VTCN1, V-set domain containing T cell activation inhibitor 1
BRCA: Breast Carcinoma
CD8: a.k.a. CD8A, cluster of differentiation 8
CK-SOX10: Cytokeratin-SRY-box transcription factor 10
COX2: a.k.a. PTGS2, Prostaglandin-endoperoxide synthase 2
ER-: Estrogen Receptor Negative
FOXP3: Forkhead Box P3
GEO: Gene Expression Omnibus
IDC: Invasive Ductal Carcinoma
IDO1: Indoleamine 2,3-dioxygenase 1
IFN: Interferon
IFNG: Interferon Gamma
IHC: Immunohistochemistry
IL1: Interleukin 1
IL10: Interleukin 10
IRB: Institutional Review Board
NOS2: Nitric Oxide Synthase 2
PD1: Programmed Cell Death 1
PDL1: Programmed Cell Death Ligand 1
TCGA: The Cancer Genome Atlas
Teff: T effector Cell
Texh: T exhausted Cell
TGF: Tumor Growth Factor
Treg: T Regulatory Cell
TME: Tumor Microenvironment
TNF: Tumor Necrosis Factor
PGE2: Prostaglandin E2
S-UMAP: Spatial Uniform Manifold Approximation and Projection

## Figure legends

**Figure Supplemental 1. Determination of significance of Thresholding of NOS2 and COX2 at the single cell level.**

The threshold was determined as previously described. A) Tables comparing one tail of Mann Whitney and Welch of the mean difference between deceased and alive. B) Ratio of NOS2 at medium and weak as well COX2 with the tumor marker CK-SOX10. C) Graphic comparison of the NOS2/tumor relationship with 2 tail MW or tail Welch. D) Linear regression analysis of the phenotype tumor-NOS2 and tumor-COX2 all samples (n=21) deceased (n=11) and alive (n=10) tumor NOS2 was used at NOS2m. The ratio of NOS2s with cellular immune and tumor cellular phenotypes where F) CD8 G) CD4 H) CD68 I) tumor. The linear regression comparing survival status J) IFNγ with NOS2s and K) IFNγ with COX2 and L) T_eff_ and NOS2.

**Supplemental Figure 2. Analysis of cellular phenotypes.**

Analysis of the ratio of cellular phenotypes comparing A) tumor-COX2 and B) NOS2s. (*p <0.5, **p <0.01, *** p<0.001, Mann-Whitney test, one-tail).

**Supplemental Figure 3. Whole tumor phenotypes.**

Defining the whole tumor phenotypes based on CD8, NOS2 and COX2 % cells. A) is the % cells distribution comparing deceased and Alive for CD8, NOS2, and COX2. B) distribution lots of tumor-NOS2 or tumor-COX2 with CD3CD8 C) table of the different phenotypes comparing deceased and alive.

## References

1. Shiravand, Y., Khodadadi, F., Kashani, S. M. A., Hosseini-Fard, S. R., Hosseini, S., Sadeghirad, H., Ladwa, R., O’Byrne, K., and Kulasinghe, A. (2022) Immune Checkpoint Inhibitors in Cancer Therapy. Curr Oncol 29, 3044–3060

2. Kleponis, J., Skelton, R., and Zheng, L. (2015) Fueling the engine and releasing the break: combinational therapy of cancer vaccines and immune checkpoint inhibitors. Cancer Biol Med 12, 201–208

3. Liu, J., Fu, M. Y., Wang, M. N., Wan, D. D., Wei, Y. Q., and Wei, X. W. (2022) Cancer vaccines as promising immuno-therapeutics: platforms and current progress. J Hematol Oncol 15

4. Vajari, M. K., Sanaei, M. J., Salari, S., Rezvani, A., Ravari, M. S., and Bashash, D. (2023) Breast cancer vaccination: Latest advances with an analytical focus on clinical trials. Int Immunopharmacol 123, 110696

5. Leowattana, W., Leowattana, P., and Leowattana, T. (2023) Immunotherapy for advanced gastric cancer. World J Methodol 13, 79–97

6. Secondino, S., Canino, C., Alaimo, D., Muzzana, M., Galli, G., Borgetto, S., Basso, S., Bagnarino, J., Pulvirenti, C., Comoli, P., and Pedrazzoli, P. (2023) Clinical Trials of Cellular Therapies in Solid Tumors. Cancers (Basel) 15

7. Valenza, C., Trapani, D., Fusco, N., Wang, X., Cristofanilli, M., Ueno, N. T., and Curigliano, G. (2023) The immunogram of inflammatory breast cancer. Cancer Treat Rev 119, 102598

8. Saleh, R., and Elkord, E. (2019) Treg-mediated acquired resistance to immune checkpoint inhibitors. Cancer Lett 457, 168–179

9. van Gulijk, M., van Krimpen, A., Schetters, S., Eterman, M., van Elsas, M., Mankor, J., Klaase, L., de Bruijn, M., van Nimwegen, M., van Tienhoven, T., van Ijcken, W., Boon, L., van der Schoot, J., Verdoes, M., Scheeren, F., van der Burg, S. H., Lambrecht, B. N., Stadhouders, R., Dammeijer, F., Aerts, J., and van Hall, T. (2023) PD-L1 checkpoint blockade promotes regulatory T cell activity that underlies therapy resistance. Sci Immunol 8

10. Anu, R. I., Shiu, K. K., and Khan, K. H. (2023) The immunomodulatory role of IDO1-Kynurenine-NAD(+) pathway in switching cold tumor microenvironment in PDAC. Front Oncol 13, 1142838

11. Stone, T. W., and Williams, R. O. (2023) Interactions of IDO and the Kynurenine Pathway with Cell Transduction Systems and Metabolism at the Inflammation-Cancer Interface. Cancers 15

12. Cheng, R. Y. S., Ridnour, L. A., Wink, A. L., Gonzalez, A. L., Femino, E. L., Rittscher, H., Somasundaram, V., Heinz, W. F., Coutinho, L., Rangel, M. C., Edmondson, E. F., Butcher, D., Kinders, R. J., Li, X. X., Wong, S. T. C., McVicar, D. W., Anderson, S. K., Pore, M., Hewitt, S. M., Billiar, T. R., Glynn, S. A., Chang, J. C., Lockett, S. J., Ambs, S., and Wink, D. A. (2023) Interferon-gamma is quintessential for NOS2 and COX2 expression in ER- breast tumors that lead to poor outcome. Cell Death Dis 14

13. Dios-Barbeito, S., Gonzalez, R., Cadenas, M., Garcia, L. F., Victor, V. M., Padillo, F. J., and Muntane, J. (2022) Impact of nitric oxide in liver cancer microenvironment. Nitric Oxide 128, 1–11

14. Liao, W., Ye, T., and Liu, H. (2019) Prognostic Value of Inducible Nitric Oxide Synthase (iNOS) in Human Cancer: A Systematic Review and Meta-Analysis. Biomed Res Int 2019, 6304851

15. Wu, M. Q., Wu, X. L., Wang, X., Hong, X. C., Liu, Y. F., Lv, G. Z., Li, C., Pan, Z. Z., Zhang, R. X., and Chen, G. (2023) IDO1/COX2 Expression Is Associated with Poor Prognosis in Colorectal Cancer Liver Oligometastases. J Pers Med 13

16. Basudhar, D., Bharadwaj, G., Somasundaram, V., Cheng, R. Y. S., Ridnour, L. A., Fujita, M., Lockett, S. J., Anderson, S. K., McVicar, D. W., and Wink, D. A. (2019) Understanding the tumour micro-environment communication network from an NOS2/COX2 perspective. Br J Pharmacol 176, 155–176

17. Somasundaram, V., Gilmore, A. C., Basudhar, D., Palmieri, E. M., Scheiblin, D. A., Heinz, W. F., Cheng, R. Y. S., Ridnour, L. A., Altan-Bonnet, G., Lockett, S. J., McVicar, D. W., and Wink, D. A. (2020) Inducible nitric oxide synthase-derived extracellular nitric oxide flux regulates proinflammatory responses at the single cell level. Redox Biology 28

18. Somasundaram, V., Basudhar, D., Bharadwaj, G., No, J. H., Ridnour, L. A., Cheng, R. Y. S., Fujita, M., Thomas, D. D., Anderson, S. K., McVicar, D. W., and Wink, D. A. (2019) Molecular Mechanisms of Nitric Oxide in Cancer Progression, Signal Transduction, and Metabolism. Antioxid Redox Signal 30, 1124–1143

19. Wang, R., Geller, D. A., Wink, D. A., Cheng, B., and Billiar, T. R. (2020) NO and hepatocellular cancer. Br J Pharmacol 177, 5459–5466

20. Lin, K., Baritaki, S., Vivarelli, S., Falzone, L., Scalisi, A., Libra, M., and Bonavida, B. (2022) The Breast Cancer Protooncogenes HER2, BRCA1 and BRCA2 and Their Regulation by the iNOS/NOS2 Axis. Antioxidants-Basel 11

21. Mohsin, N. U., Aslam, S., Ahmad, M., Irfan, M., Al-Hussain, S. A., and Zaki, M. E. A. (2022) Cyclooxygenase-2 (COX-2) as a Target of Anticancer Agents: A Review of Novel Synthesized Scaffolds Having Anticancer and COX-2 Inhibitory Potentialities. Pharmaceuticals-Base 15

22. Jin, K. P., Qian, C., Lin, J. T., and Liu, B. (2023) Cyclooxygenase-2-Prostaglandin E2 pathway: A key player in tumor-associated immune cells. Frontiers in Oncology 13

23. Sahu, A., Raza, K., Pradhan, D., Jain, A. K., and Verma, S. (2023) Cyclooxygenase-2 as a therapeutic target against human breast cancer: A comprehensive review. Wires Mech Dis

24. Somasundaram, V., Ridnour, L. A., Cheng, R. Y., Walke, A. J., Kedei, N., Bhattacharyya, D. D., Wink, A. L., Edmondson, E. F., Butcher, D., Warner, A. C., Dorsey, T. H., Scheiblin, D. A., Heinz, W., Bryant, R. J., Kinders, R. J., Lipkowitz, S., Wong, S. T., Pore, M., Hewitt, S. M., McVicar, D. W., Anderson, S. K., Chang, J., Glynn, S. A., Ambs, S., Lockett, S. J., and Wink, D. A. (2022) Systemic Nos2 Depletion and Cox inhibition limits TNBC disease progression and alters lymphoid cell spatial orientation and density. Redox Biol 58, 102529

25. Basudhar, D., Glynn, S. A., Greer, M., Somasundaram, V., No, J. H., Scheiblin, D. A., Garrido, P., Heinz, W. F., Ryan, A. E., Weiss, J. M., Cheng, R. Y. S., Ridnour, L. A., Lockett, S. J., McVicar, D. W., Ambs, S., and Wink, D. A. (2017) Coexpression of NOS2 and COX2 accelerates tumor growth and reduces survival in estrogen receptor-negative breast cancer. P Natl Acad Sci USA 114, 13030–13035

26. Davila-Gonzalez, D., Chang, J. C., and Billiar, T. R. (2017) NO and COX2: Dual targeting for aggressive cancers. P Natl Acad Sci USA 114, 13591–13593

27. 27. Landskron, G., De la Fuente, M., Thuwajit, P., Thuwajit, C., and Hermoso, M. A. (2014) Chronic Inflammation and Cytokines in the Tumor Microenvironment. J Immunol Res 2014

28. McGinity, C. L., Palmieri, E. M., Somasundaram, V., Bhattacharyya, D. D., Ridnour, L. A., Cheng, R. Y. S., Ryan, A. E., Glynn, S. A., Thomas, D. D., Miranda, K. M., Anderson, S. K., Lockett, S. J., McVicar, D. W., and Wink, D. A. (2021) Nitric Oxide Modulates Metabolic Processes in the Tumor Immune Microenvironment. Int J Mol Sci 22

29. Ladetto, M., Vallet, S., Trojan, A., Dell’Aquila, M., Monitillo, L., Rosato, R., Santo, L., Drandi, D., Bertola, A., Falco, P., Cavallo, F., Ricca, I., De Marco, F., Mantoan, B., Bode-Lesniewska, B., Pagliano, G., Francese, R., Rocci, A., Astolfi, M., Compagno, M., Mariani, S., Godio, L., Marino, L., Ruggeri, M., Omedè, P., Palumbo, A., and Boccadoro, M. (2005) Cyclooxygenase-2 (COX-2) is frequently expressed in multiple myeloma and is an independent predictor of poor outcome. Blood 105, 4784–4791

30. Liu, Y., Xu, R. Y., Gu, H. Y., Zhang, E. F., Qu, J. W., Cao, W., Huang, X., Yan, H. M., He, J. S., and Cai, Z. (2021) Metabolic reprogramming in macrophage responses. Biomark Res 9

31. Salvemini, D., Kim, S. F., and Mollace, V. (2013) Reciprocal regulation of the nitric oxide and cyclooxygenase pathway in pathophysiology: relevance and clinical implications. Am J Physiol-Reg I 304, R473–R487

32. Prendergast, G. C., Mondal, A., Dey, S., Laury-Kleintop, L. D., and Muller, A. J. (2018) Inflammatory Reprogramming with IDO1 Inhibitors: Turning Immunologically Unresponsive ’Cold’ Tumors ’Hot’. Trends Cancer 4, 38–58

33. Janssen, L. M. E., Ramsay, E. E., Logsdon, C. D., and Overwijk, W. W. (2017) The immune system in cancer metastasis: friend or foe? J Immunother Cancer 5, 79

34. Jung, M. Y., Aibaidula, A., Brown, D. A., Himes, B. T., Garcia, L. M. C., and Parney, I. F. (2022) Superinduction of immunosuppressive glioblastoma extracellular vesicles by IFN-γ through PD-L1 and IDO1. Neuro-Oncol Adv 4

35. Garcia-Diaz, A., Shin, D. S., Moreno, B. H., Saco, J., Escuin-Ordinas, H., Rodriguez, G. A., Zaretsky, J. M., Sun, L., Hugo, W., Wang, X. Y., Parisi, G., Saus, C. P., Torrejon, D. Y., Graeber, T. G., Comin-Anduix, B., Hu-Lieskovan, S., Damoiseaux, R., Lo, R. S., and Ribas, A. (2019) Interferon Receptor Signaling Pathways Regulating PD-L1 and PD-L2 Expression (vol 19, pg 1189, 2017). Cell Rep 29, 3766–3766

36. Zhu, Y., Zhao, Y., Cao, Z., Chen, Z., and Pan, W. (2022) Identification of three immune subtypes characterized by distinct tumor immune microenvironment and therapeutic response in stomach adenocarcinoma. Gene 818, 146177

37. Fu, T., Dai, L. J., Wu, S. Y., Xiao, Y., Ma, D., Jiang, Y. Z., and Shao, Z. M. (2021) Spatial architecture of the immune microenvironment orchestrates tumor immunity and therapeutic response. J Hematol Oncol 14, 98

38. Wu, T., and Dai, Y. (2017) Tumor microenvironment and therapeutic response. Cancer Lett 387, 61–68

39. Barnestein, R., Galland, L., Kalfeist, L., Ghiringhelli, F., Ladoire, S., and Limagne, E. (2022) Immunosuppressive tumor microenvironment modulation by chemotherapies and targeted therapies to enhance immunotherapy effectiveness. Oncoimmunology 11

40. Schalper, K. A., Carvajal-Hausdorf, D., McLaughlin, J., Altan, M., Velcheti, V., Gaule, P., Sanmamed, M. F., Chen, L., Herbst, R. S., and Rimm, D. L. (2017) Differential Expression and Significance of PD-L1, IDO-1, and B7-H4 in Human Lung Cancer. Clin Cancer Res 23, 370–378

41. Zou, W. P., Wolchok, J. D., and Chen, L. P. (2016) PD-L1 (B7-H1) and PD-1 pathway blockade for cancer therapy: Mechanisms, response biomarkers, and combinations. Sci Transl Med 8

42. Komai, T., Inoue, M., Okamura, T., Morita, K., Iwasaki, Y., Sumitomo, S., Shoda, H., Yamamoto, K., and Fujio, K. (2018) Transforming Growth Factor-β and Interleukin-10 Synergistically Regulate Humoral Immunity Modulating Metabolic Signals. Front Immunol 9

43. Thepmalee, C., Panya, A., Junking, M., Chieochansin, T., and Yenchitsomanus, P. T. (2018) Inhibition of IL-10 and TGF-beta receptors on dendritic cells enhances activation of effector T-cells to kill cholangiocarcinoma cells. Hum Vaccin Immunother 14, 1423–1431

44. Routy, J. P., Routy, B., Graziani, G. M., and Mehraj, V. (2016) The Kynurenine Pathway Is a Double-Edged Sword in Immune-Privileged Sites and in Cancer: Implications for Immunotherapy. Int J Tryptophan Res 9, 67–77

45. Lian, J. C., Liang, Y. F., Zhang, H. L., Lan, M. S., Ye, Z. Y., Lin, B. H., Qiu, X. X., and Zeng, J. C. (2022) The role of polyamine metabolism in remodeling immune responses and blocking therapy within the tumor immune microenvironment. Front Immunol 13

46. Chia, T. Y., Zolp, A., and Miska, J. (2022) Polyamine Immunometabolism: Central Regulators of Inflammation, Cancer and Autoimmunity. Cells-Basel 11

47. Wang, D., and DuBois, R. N. (2016) The Role of Prostaglandin E(2) in Tumor-Associated Immunosuppression. Trends Mol Med 22, 1–3

48. Ekmekcioglu, S., Grimm, E. A., and Roszik, J. (2017) Targeting iNOS to increase efficacy of immunotherapies. Hum Vaccin Immunother 13, 1105–1108

49. Glynn, S. A., Boersma, B. J., Dorsey, T. H., Yi, M., Yfantis, H. G., Ridnour, L. A., Martin, D. N., Switzer, C. H., Hudson, R. S., Wink, D. A., Lee, D. H., Stephens, R. M., and Ambs, S. (2010) Increased NOS2 predicts poor survival in estrogen receptor-negative breast cancer patients. J Clin Invest 120, 3843–3854

50. Glynn, S. A., Prueitt, R. L., Ridnour, L. A., Boersma, B. J., Dorsey, T. M., Wink, D. A., Goodman, J. E., Yfantis, H. G., Lee, D. H., and Ambs, S. (2010) COX-2 activation is associated with Akt phosphorylation and poor survival in ER-negative, HER2-positive breast cancer. BMC Cancer 10, 626

51. Jorgovanovic, D., Song, M. J., Wang, L. P., and Zhang, Y. (2020) Roles of IFN-γ in tumor progression and regression: a review. Biomark Res 8

52. Enayati, S., Seifirad, S., Amiri, P., Abolhalaj, M., and Mohammad-Amoli, M. (2015) Interleukin-1 beta, interferon-gamma, and tumor necrosis factor-alpha gene expression in peripheral blood mononuclear cells of patients with coronary artery disease. Arya Atheroscler 11, 267–274

53. Kartikasari, A. E. R., Huertas, C. S., Mitchell, A., and Plebanski, M. (2021) Tumor-Induced Inflammatory Cytokines and the Emerging Diagnostic Devices for Cancer Detection and Prognosis. Frontiers in Oncology 11

54. Lenicov, F. R., Paletta, A. L., Prinz, M. G., Varese, A., Pavillet, C. E., Malizia, A. L., Sabatté, J., Geffner, J. R., and Ceballos, A. (2018) Prostaglandin E2 Antagonizes TGF-β Actions During the Differentiation of Monocytes Into Dendritic Cells. Front Immunol 9

55. MacKenzie, K. F., Clark, K., Naqvi, S., McGuire, V. A., Noehren, G., Kristariyanto, Y., van den Bosch, M., Mudaliar, M., McCarthy, P. C., Pattison, M. J., Pedrioli, P. G., Barton, G. J., Toth, R., Prescott, A., and Arthur, J. S. (2013) PGE(2) induces macrophage IL-10 production and a regulatory-like phenotype via a protein kinase A-SIK-CRTC3 pathway. J Immunol 190, 565–577

56. Gruosso, T., Gigoux, M., Manem, V. S. K., Bertos, N., Zuo, D., Perlitch, I., Saleh, S. M. I., Zhao, H., Souleimanova, M., Johnson, R. M., Monette, A., Ramos, V. M., Hallett, M. T., Stagg, J., Lapointe, R., Omeroglu, A., Meterissian, S., Buisseret, L., Van den Eynden, G., Salgado, R., Guiot, M. C., Haibe-Kains, B., and Park, M. (2019) Spatially distinct tumor immune microenvironments stratify triple-negative breast cancers. J Clin Invest 129, 1785–1800

57. Oyler-Yaniv, J., Oyler-Yaniv, A., Shakiba, M., Min, N. K., Chen, Y. H., Cheng, S. Y., Krichevsky, O., Altan-Bonnet, N., and Altan-Bonnet, G. (2017) Catch and Release of Cytokines Mediated by Tumor Phosphatidylserine Converts Transient Exposure into Long-Lived Inflammation. Mol Cell 66, 635–647 e637

58. Giraldo, N. A., Berry, S., Becht, E., Ates, D., Schenk, K. M., Engle, E. L., Green, B., Nguyen, P., Soni, A., Stein, J. E., Succaria, F., Ogurtsova, A., Xu, H., Gottardo, R., Anders, R. A., Lipson, E. J., Danilova, L., Baras, A. S., and Taube, J. M. (2021) Spatial UMAP and Image Cytometry for Topographic Immuno-oncology Biomarker Discovery. Cancer Immunol Res 9, 1262–1269

59. Altan, M., Kidwell, K. M., Pelekanou, V., Carvajal-Hausdorf, D. E., Schalper, K. A., Toki, M. I., Thomas, D. G., Sabel, M. S., Hayes, D. F., and Rimm, D. L. (2018) Association of B7-H4, PD-L1, and tumor infiltrating lymphocytes with outcomes in breast cancer. NPJ Breast Cancer 4, 40

60. Xu, Y., Zhu, S., Song, M., Liu, W., Liu, C., Li, Y., and Wang, M. (2014) B7-H4 expression and its role in interleukin-2/interferon treatment of clear cell renal cell carcinoma. Oncol Lett 7, 1474–1478

61. Soliman, H., Rawal, B., Fulp, J., Lee, J. H., Lopez, A., Bui, M. M., Khalil, F., Antonia, S., Yfantis, H. G., Lee, D. H., Dorsey, T. H., and Ambs, S. (2013) Analysis of indoleamine 2-3 dioxygenase (IDO1) expression in breast cancer tissue by immunohistochemistry. Cancer Immunol Immunother 62, 829–837

62. Jiang, W., He, Y., He, W., Wu, G., Zhou, X., Sheng, Q., Zhong, W., Lu, Y., Ding, Y., Lu, Q., Ye, F., and Hua, H. (2020) Exhausted CD8+T Cells in the Tumor Immune Microenvironment: New Pathways to Therapy. Front Immunol 11, 622509

63. Pluhar, G. E., Pennell, C. A., and Olin, M. R. (2015) CD8(+) T Cell-Independent Immune-Mediated Mechanisms of Anti-Tumor Activity. Crit Rev Immunol 35, 153–172

64. Mortezaee, K., and Majidpoor, J. (2023) Mechanisms of CD8(+) T cell exclusion and dysfunction in cancer resistance to anti-PD-(L)1. Biomed Pharmacother 163, 114824

65. Trefny, M. P., Kirchhammer, N., Auf der Maur, P., Natoli, M., Schmid, D., Germann, M., Fernandez Rodriguez, L., Herzig, P., Lotscher, J., Akrami, M., Stinchcombe, J. C., Stanczak, M. A., Zingg, A., Buchi, M., Roux, J., Marone, R., Don, L., Lardinois, D., Wiese, M., Jeker, L. T., Bentires-Alj, M., Rossy, J., Thommen, D. S., Griffiths, G. M., Laubli, H., Hess, C., and Zippelius, A. (2023) Deletion of SNX9 alleviates CD8 T cell exhaustion for effective cellular cancer immunotherapy. Nat Commun 14, 86

66. Oshi, M., Asaoka, M., Tokumaru, Y., Yan, L., Matsuyama, R., Ishikawa, T., Endo, I., and Takabe, K. (2020) CD8 T Cell Score as a Prognostic Biomarker for Triple Negative Breast Cancer. Int J Mol Sci 21

67. Miyashita, M., Sasano, H., Tamaki, K., Hirakawa, H., Takahashi, Y., Nakagawa, S., Watanabe, G., Tada, H., Suzuki, A., Ohuchi, N., and Ishida, T. (2015) Prognostic significance of tumor-infiltrating CD8+ and FOXP3+ lymphocytes in residual tumors and alterations in these parameters after neoadjuvant chemotherapy in triple-negative breast cancer: a retrospective multicenter study. Breast Cancer Res 17, 124

68. Huertas-Caro, C. A., Ramirez, M. A., Rey-Vargas, L., Bejarano-Rivera, L. M., Ballen, D. F., Nunez, M., Mejia, J. C., Sua-Villegas, L. F., Cock-Rada, A., Zabaleta, J., Fejerman, L., Sanabria-Salas, M. C., and Serrano-Gomez, S. J. (2023) Tumor infiltrating lymphocytes (TILs) are a prognosis biomarker in Colombian patients with triple negative breast cancer. Sci Rep 13, 21324

69. Li, F., Li, C. C., Cai, X. Y., Xie, Z. H., Zhou, L. Q., Cheng, B., Zhong, R., Xiong, S., Li, J. F., Chen, Z. X., Yu, Z. W., He, J. X., and Liang, W. H. (2021) The association between CD8+tumor-infiltrating lymphocytes and the clinical outcome of cancer immunotherapy: A systematic review and meta-analysis. Eclinicalmedicine 41

70. Fridman, W. H., Pages, F., Sautes-Fridman, C., and Galon, J. (2012) The immune contexture in human tumours: impact on clinical outcome. Nat Rev Cancer 12, 298–306

71. Saito, T., Nishikawa, H., Wada, H., Nagano, Y., Sugiyama, D., Atarashi, K., Maeda, Y., Hamaguchi, M., Ohkura, N., Sato, E., Nagase, H., Nishimura, J., Yamamoto, H., Takiguchi, S., Tanoue, T., Suda, W., Morita, H., Hattori, M., Honda, K., Mori, M., Doki, Y., and Sakaguchi, S. (2016) Two FOXP3(+)CD4(+) T cell subpopulations distinctly control the prognosis of colorectal cancers. Nat Med 22, 679–684

72. Sautes-Fridman, C., Petitprez, F., Calderaro, J., and Fridman, W. H. (2019) Tertiary lymphoid structures in the era of cancer immunotherapy. Nat Rev Cancer 19, 307–325

73. Wan, Y. Y., and Flavell, R. A. (2007) ’Yin-Yang’ functions of transforming growth factor-beta and T regulatory cells in immune regulation. Immunol Rev 220, 199–213

74. Chen, W. J. (2023) TGF-β Regulation of T Cells. Annu Rev Immunol 41, 483–512

75. Chapoval, S., Dasgupta, P., Dorsey, N. J., and Keegan, A. D. (2010) Regulation of the T helper cell type 2 (Th2)/T regulatory cell (Treg) balance by IL-4 and STAT6. J Leukoc Biol 87, 1011–1018

76. Ju, X., Zhang, H., Zhou, Z., Chen, M., and Wang, Q. (2020) Tumor-associated macrophages induce PD-L1 expression in gastric cancer cells through IL-6 and TNF-a signaling. Exp Cell Res 396, 112315

77. Zhang, H., Liu, L., Liu, J. B., Dang, P. Y., Hu, S. Y., Yuan, W. T., Sun, Z. Q., Liu, Y., and Wang, C. Z. (2023) Roles of tumor-associated macrophages in anti-PD-1/PD-L1 immunotherapy for solid cancers. Mol Cancer 22

78. Antonangeli, F., Natalini, A., Garassino, M. C., Sica, A., Santoni, A., and Di Rosa, F. (2020) Regulation of PD-L1 Expression by NF-κB in Cancer. Front Immunol 11

79. Cavalcante, R. S., Ishikawa, U., Silva, E. S., Silva-Junior, A. A., Araujo, A. A., Cruz, L. J., Chan, A. B., and de Araujo Junior, R. F. (2021) STAT3/NF-kappaB signalling disruption in M2 tumour-associated macrophages is a major target of PLGA nanocarriers/PD-L1 antibody immunomodulatory therapy in breast cancer. Br J Pharmacol 178, 2284–2304

80. Pu, Y., and Ji, Q. (2022) Tumor-Associated Macrophages Regulate PD-1/PD-L1 Immunosuppression. Front Immunol 13, 874589

81. Cai, H., Zhang, Y. C., Wang, J., and Gu, J. Y. (2021) Defects in Macrophage Reprogramming in Cancer Therapy: The Negative Impact of PD-L1/PD-1. Front Immunol 12

82. Quaranta, V., Rainer, C., Nielsen, S. R., Raymant, M. L., Ahmed, M. S., Engle, D. D., Taylor, A., Murray, T., Campbell, F., Palmer, D. H., Tuveson, D. A., Mielgo, A., and Schmid, M. C. (2018) Macrophage-Derived Granulin Drives Resistance to Immune Checkpoint Inhibition in Metastatic Pancreatic Cancer. Cancer Res 78, 4253–4269

83. Spranger, S., Spaapen, R. M., Zha, Y., Williams, J., Meng, Y., Ha, T. T., and Gajewski, T. F. (2013) Up-Regulation of PD-L1, IDO, and T(reg) in the Melanoma Tumor microenvironment Is Driven by CD8(+) T Cells. Sci Transl Med 5

84. Liu, X. W., Yan, G. Y., Xu, B. Y., Yu, H., An, Y., and Sun, M. J. (2022) Evaluating the role of IDO1 macrophages in immunotherapy using scRNA-seq and bulk-seq in colorectal cancer. Front Immunol 13

85. Toulmonde, M., Penel, N., Adam, J., Chevreau, C., Blay, J. Y., Le Cesne, A., Bompas, E., Piperno-Neumann, S., Cousin, S., Grellety, T., Ryckewaert, T., Bessede, A., Ghiringhelli, F., Pulido, M., and Italiano, A. (2018) Use of PD-1 Targeting, Macrophage Infiltration, and IDO Pathway Activation in Sarcomas A Phase 2 Clinical Trial. Jama Oncol 4, 93–97

86. Wink, D. A., Hines, H. B., Cheng, R. Y., Switzer, C. H., Flores-Santana, W., Vitek, M. P., Ridnour, L. A., and Colton, C. A. (2011) Nitric oxide and redox mechanisms in the immune response. J Leukoc Biol 89, 873–891

87. Bonaventura, P., Shekarian, T., Alcazer, V., Valladeau-Guilemond, J., Valsesia-Wittmann, S., Amigorena, S., Caux, C., and Depil, S. (2019) Cold Tumors: A Therapeutic Challenge for Immunotherapy. Front Immunol 10, 168

88. Lin, B. S., Du, L. K., Li, H. M., Zhu, X., Cui, L., and Li, X. S. (2020) Tumor-infiltrating lymphocytes: Warriors fight against tumors powerfully. Biomedicine & Pharmacotherapy 132

89. Kazemi, M. H., Sadri, M., Najafi, A., Rahimi, A., Baghernejadan, Z., Khorramdelazad, H., and Falak, R. (2022) Tumor-infiltrating lymphocytes for treatment of solid tumors: It takes two to tango? Front Immunol 13, 1018962

90. Brummel, K., Eerkens, A. L., de Bruyn, M., and Nijman, H. W. (2023) Tumour-infiltrating lymphocytes: from prognosis to treatment selection. Brit J Cancer 128, 451–458

91. Chung, A. W., Anand, K., Anselme, A. C., Chan, A. A., Gupta, N., Venta, L. A., Schwartz, M. R., Qian, W., Xu, Y., Zhang, L., Kuhn, J., Patel, T., Rodriguez, A. A., Belcheva, A., Darcourt, J., Ensor, J., Bernicker, E., Pan, P. Y., Chen, S. H., Lee, D. J., Niravath, P. A., and Chang, J. C. (2021) A phase 1/2 clinical trial of the nitric oxide synthase inhibitor L-NMMA and taxane for treating chemoresistant triple-negative breast cancer. Sci Transl Med 13, eabj5070

92. Kwa, M. J., and Adams, S. (2018) Checkpoint Inhibitors in Triple-Negative Breast Cancer (TNBC): Where to Go From Here. Cancer-Am Cancer Soc 124, 2086–2103

93. Jungles, K. M., Holcomb, E. A., Pearson, A. N., Jungles, K. R., Bishop, C. R., Pierce, L. J., Green, M. D., and Speers, C. W. (2022) Updates in combined approaches of radiotherapy and immune checkpoint inhibitors for the treatment of breast cancer. Frontiers in Oncology 12

94. Manesse, M., Patel, K. K., Bobrow, M., and Downing, S. R. (2020) The InSituPlex((R)) Staining Method for Multiplexed Immunofluorescence Cell Phenotyping and Spatial Profiling of Tumor FFPE Samples. Methods Mol Biol 2055, 585–592

