## Supplementary Figures for "Spatial analysis of NOS2 and COX2 interaction with T-effector cells reveals immunosuppressive landscapes associated with poor outcome in ER- breast cancer patients"

Supplement Figure 1.

This is compliment for Figure 1

A

Mann Whitney one tail analysis

| Threshold | deceased | Alive | P vlue (MW) |
|---|---|---|---|
| NOS2all | 14.2 | 6.36 | 0.049 |
| NOS2s | 1.18 | 0.32 | 0.006 |
| NOS2m | 3.1 | 1.2 | 0.0215 |
| NOS2w | 8.5 | 4.8 | 0.075 |
| COX2all | 9.5 | 6.3 | 0.0423 |
| COX2s | 0.5583 | 0.2856 | 0.2131 |
| COX2m | 1.89 | 1.028 | 0.086 |
| COX2w | 8.251 | 4.574 | 0.033 |

Welch t-test one tail

| | deceased | Alive | p value (Welch)# |
|---|---|---|---|
| NOS2all | 16.9 | 6.738 | 0.0195 |
| NOS2s | 2.9 | 0.6 | 0.025 |
| NOS2m | 4.2 | 1.14 | 0.019 |
| NOS2w | 9.7 | 4.7 | 0.025 |
| COX2all | 14.69 | 7.547 | 0.075 |
| COX2s | 1.6 | 0.4315 | 0.15 |
| COX2m | 3.102 | 1.412 | 0.12 |
| COX2w | 9.992 | 5.704 | 0.046 |

B

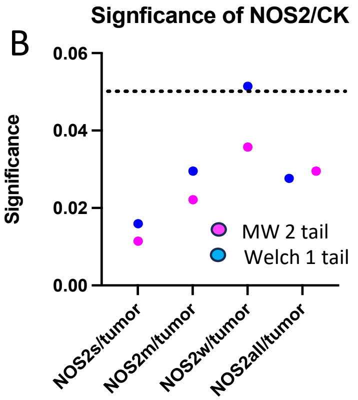


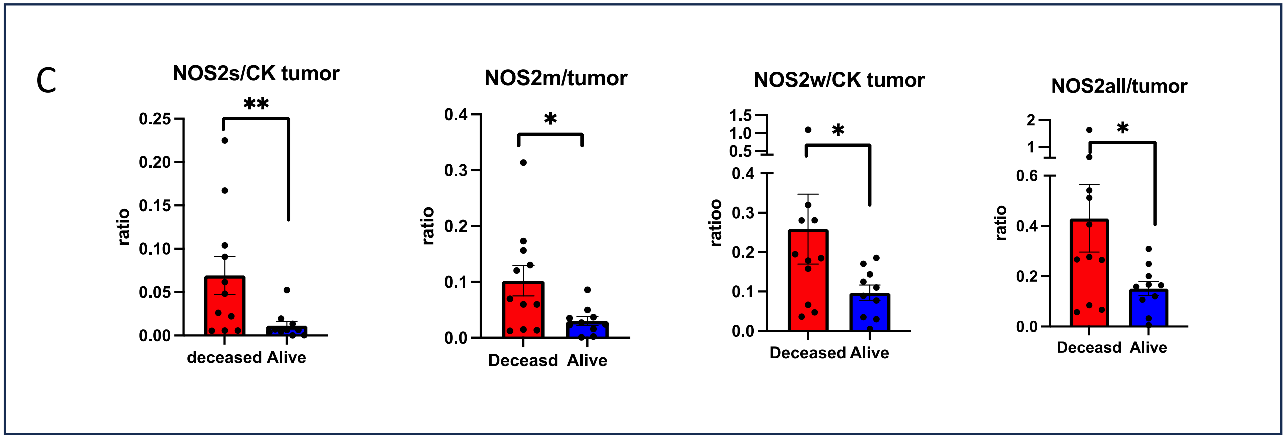


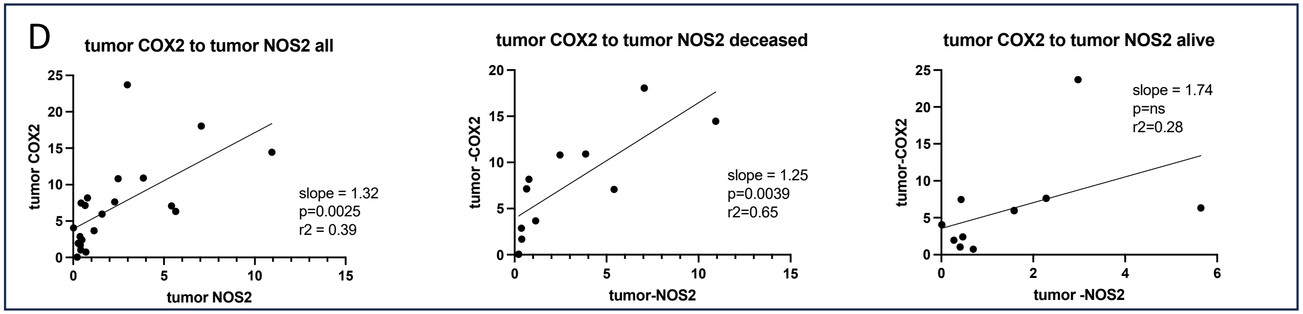


Supplemental Figure 1

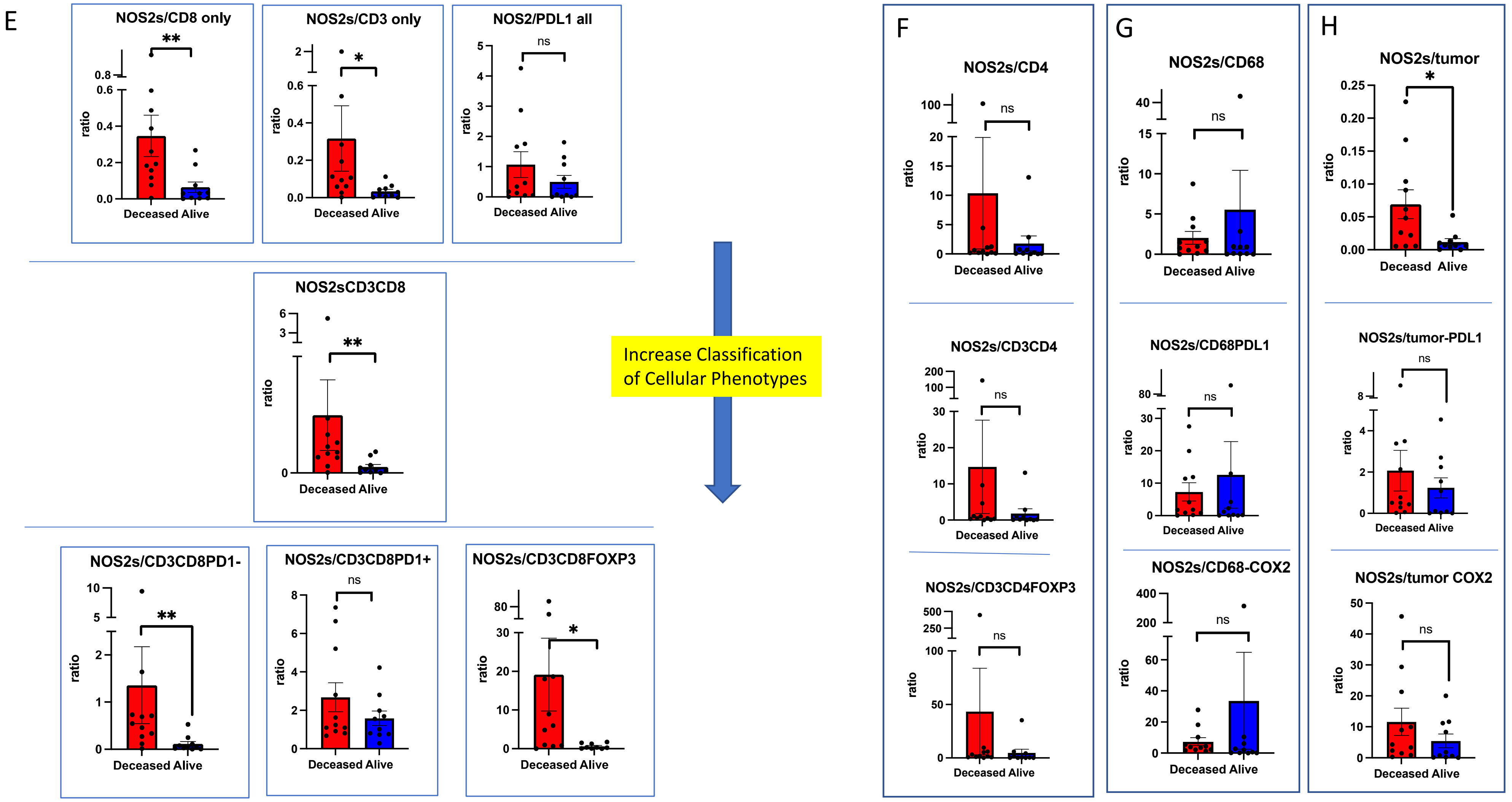


Supplement Figure 1

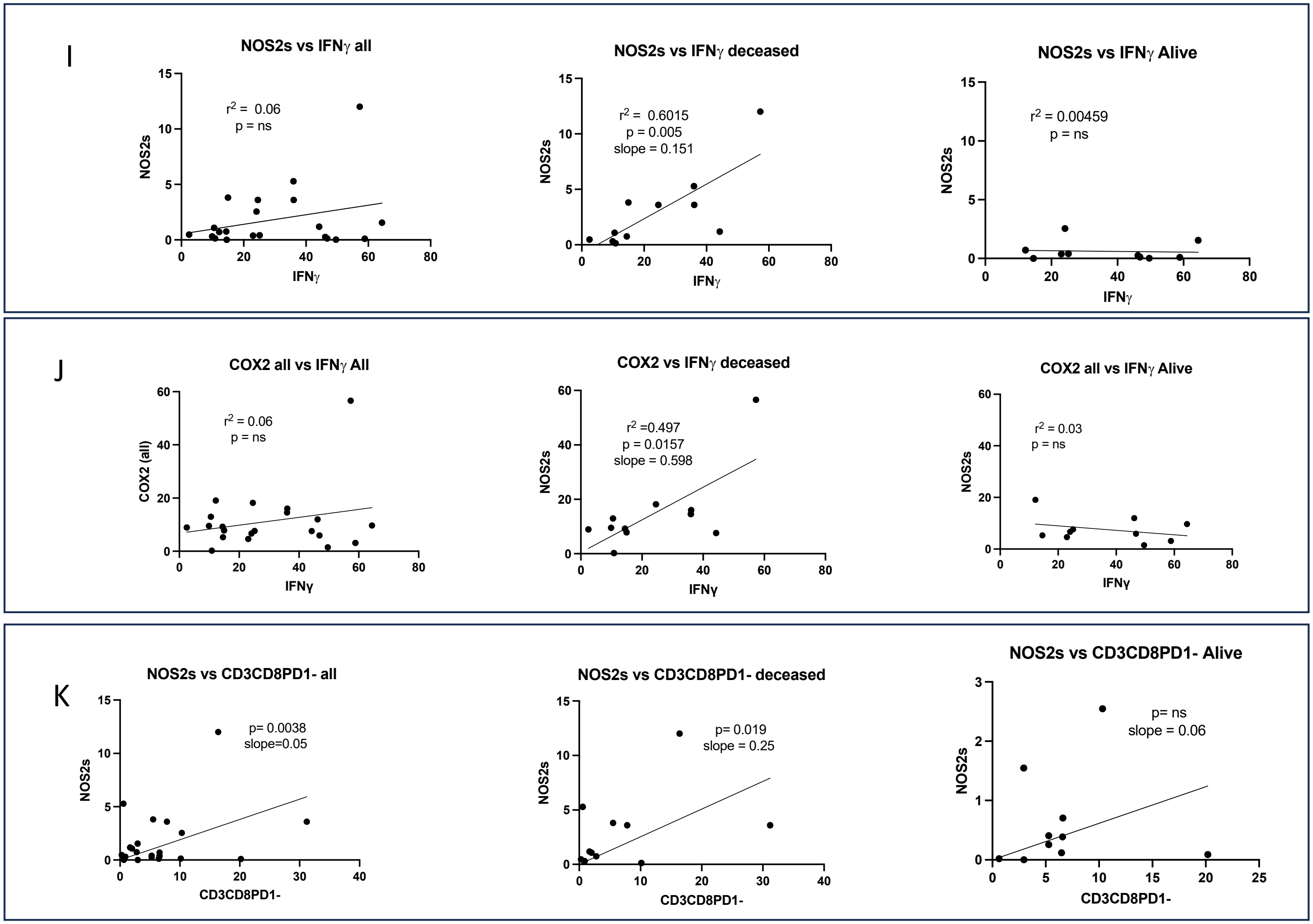


Supplemental Figure 1

Supplement Figure 2.

This is compliment for Figure 2

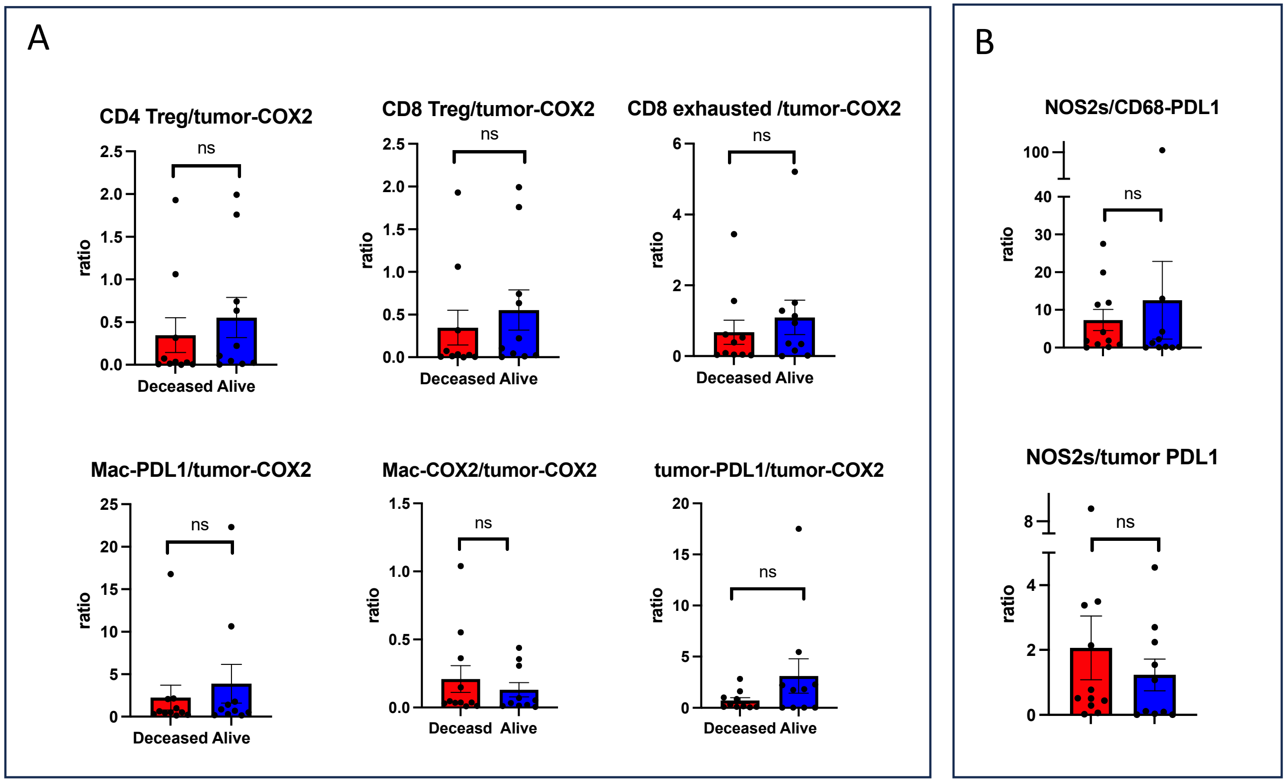


Supplement Figure 2

Supplement Figure 3.

This is compliment for Figure 5.

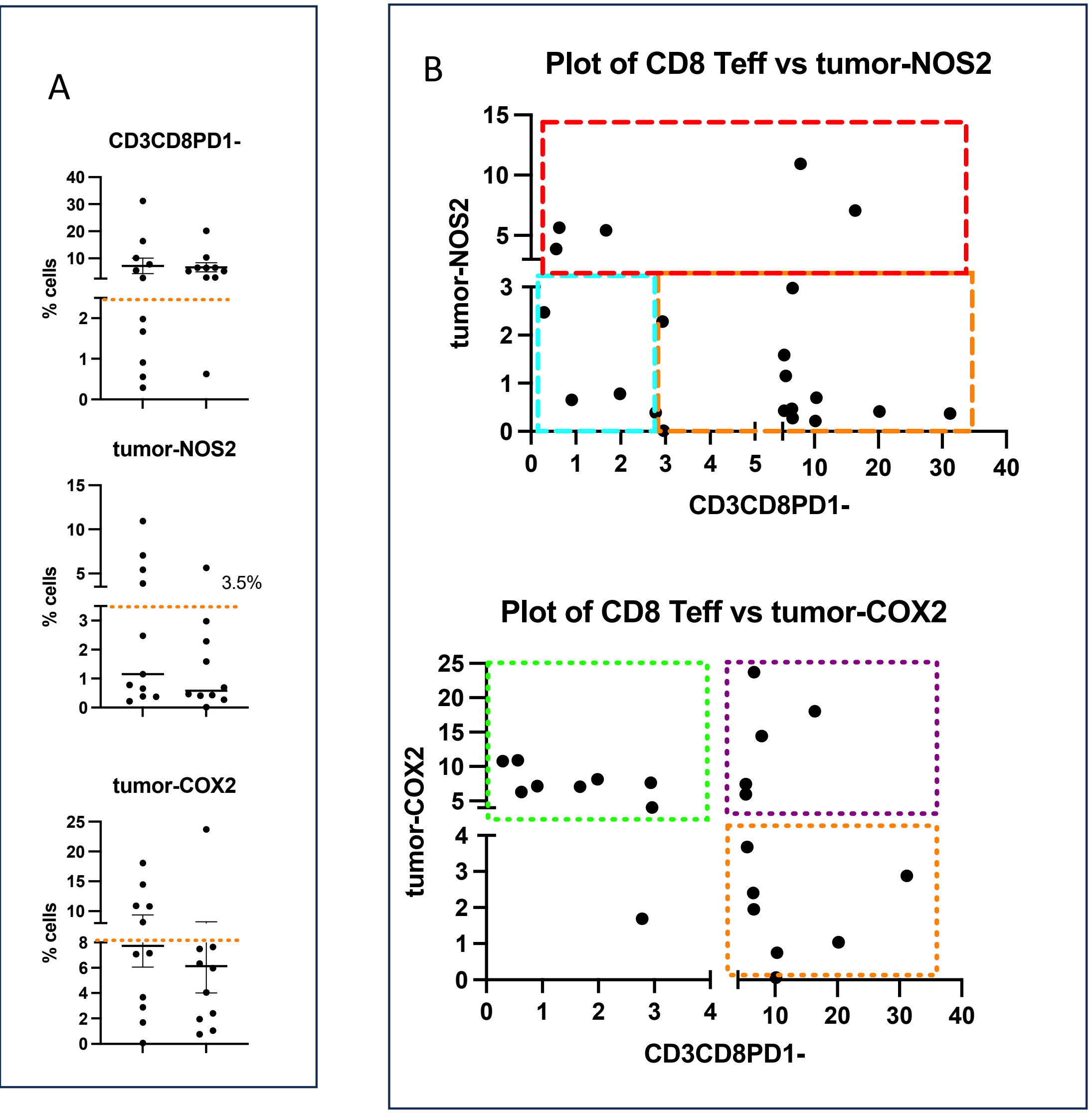


C

| | deceased | Alive |
|---|---|---|
| CD8+NOS2+COX2+ | 1 | 0 |
| CD8-NOS2-COX2+ | 6 | 1 |
| CD8+NOS2-COX2- | 3 | 9 |
| CD8-NOS2-COX2- | 1 | 0 |

Supplement figure 5

Supplemental in showing that radiation leads to marginal and increase in immune cells but no penetration
